# EFA6B regulates a stop signal for collective invasion in breast cancer

**DOI:** 10.1101/2020.03.19.998575

**Authors:** Fayad Racha, Vázquez Rojas Monserrat, Partisani Mariagrazia, Finetti Pascal, Dib Shiraz, Virolle Virginie, Cabaud Olivier, Lopez Marc, Birnbaum Daniel, Bertucci François, Franco Michel, Luton Frédéric

## Abstract

Cancer is initiated by somatic mutations in oncogenes or tumor suppressor genes, however additional mutations provide selective advantages to the tumor cells to resist treatment and develop metastases, therefore identification of secondary mutations is of paramount importance. EFA6B (Exchange Factor for ARF6, B) expression is reduced in breast cancer. To study the pro-tumoral impact of the loss of EFA6B we have invalidated its gene in normal human mammary cells. We found that EFA6B knock-out triggers a transcriptional reprogramming of the cell-to-ECM interaction machinery and unleashes CDC42-dependent collective invasion in collagen. In addition, invasive and metastatic tumors isolated from patients have lower expression of EFA6B and display gene ontology signatures identical to those of EFA6B knock-out cells. Thus, we reveal a new EFA6B-regulated molecular mechanism that controls the invasive potential of mammary cells; this finding opens up new avenues for the treatment of invasive breast cancer.

## Introduction

Breast cancer (BC) is a major public health issue with half a million deaths worldwide each year, essentially due to metastatic dissemination^1^. Despite extensive research, there is still no marker to predict the transition from *in-situ* to invasive carcinoma, and treatment of metastasis remains a largely unresolved problem. There is therefore an urgent need to understand better the molecular mechanisms that support tumor invasion.

Physiological or tumor invasion of cells within the extracellular matrix (ECM) is defined according to several criteria: individual or collective migration, non-proteolytic or degradative invasion of matrix, and structural remodeling of the extracellular matrix (ECM) to facilitate migration by applying cell-generated forces at adhesion sites^2^. It has been proposed that invasion of tumor cells is due to a lack of responsiveness to stop signals provided by the extracellular matrix (ECM). A large number of studies have described the cellular and molecular mechanisms that promote or sustain invasion but much less is known about intracellular signaling pathways that restrain the invasive potential of normal epithelial cells or transduce stop signals to neoplastic cells^3–6^.

We have previously reported the anti-tumor potential of the ARF6 exchange factor, EFA6B. We showed that its level of expression determines the epithelial status of mammary cells grown in 3D culture. In particular, the over-expression of EFA6B in weakly tumor cells restores a normal epithelial phenotype by stimulating the formation of a central lumen and tight junctions^7,8^. In BC patients we observed a correlation of the loss of expression of EFA6B with the metastatic Claudin-Low subtype and with a reduced survival rate in early BC^7^. Here, to determine the mechanism by which the loss of EFA6B might facilitate the progression of metastatic tumors, we analyzed the consequence of invalidating its gene (*PSD4*) in normal human mammary cells.

We found that EFA6B knock-out (KO) mammary cells undergo collective invasion when grown in 3D-collagen I. This invasion is supported by the activation of an epithelial-to-mesenchymal transition (EMT) program and the alteration of cell-extracellular matrix (ECM) interaction. Elucidation of the molecular mode of action reveals that EFA6B KO leads to the activation of CDC42, which in turn elicits two signaling pathways: Cdc42-MRCK-pMLC, which regulates contractility, and Cdc42-NWASP-Arp2/3, which is required for the formation of integrin β1-based and MMP14-enriched invadopodia. These two pathways are essential for invading a 3D-collagen matrix. Consistently, the expression of EFA6B is lower in invasive than in in-situ tumors isolated from patients, and the invasive tumors display gene ontology signatures identical to those of EFA6B knock-out cells. Thus, we reveal a new EFA6B-regulated molecular mechanism that controls the invasive potential of mammary cells; this finding opens up new avenues for the treatment of invasive BC.

## Results

### Loss of EFA6B in MCF10A stimulates invasive branching in collagen I

We have inactivated EFA6B gene *PSD4* in the normal human mammary cell line MCF10A using the CRISPR/Cas9 technology. **Fig.1a** shows the characterization of three homozygous (KO55, KO50 and KO2) and one heterozygous (Het2.9) KO clones, with the latter expressing half of the total levels of EFA6B compared to wild-type (WT) cells. A slight decrease (1.4±0.4 fold) of ARF6 expression was observed (**Fig.1b**) which was also noticed in BC patients whose EFA6B expression was decreased^7^. Notably, ARF6GTP levels were severely reduced (2.5±0.4 fold) indicating that EFA6B is a major ARF6-GEF in MCF10A cells (**Fig.1b**). The levels of the other EFA6 and ARF proteins remained unaffected.

**Fig.1:**
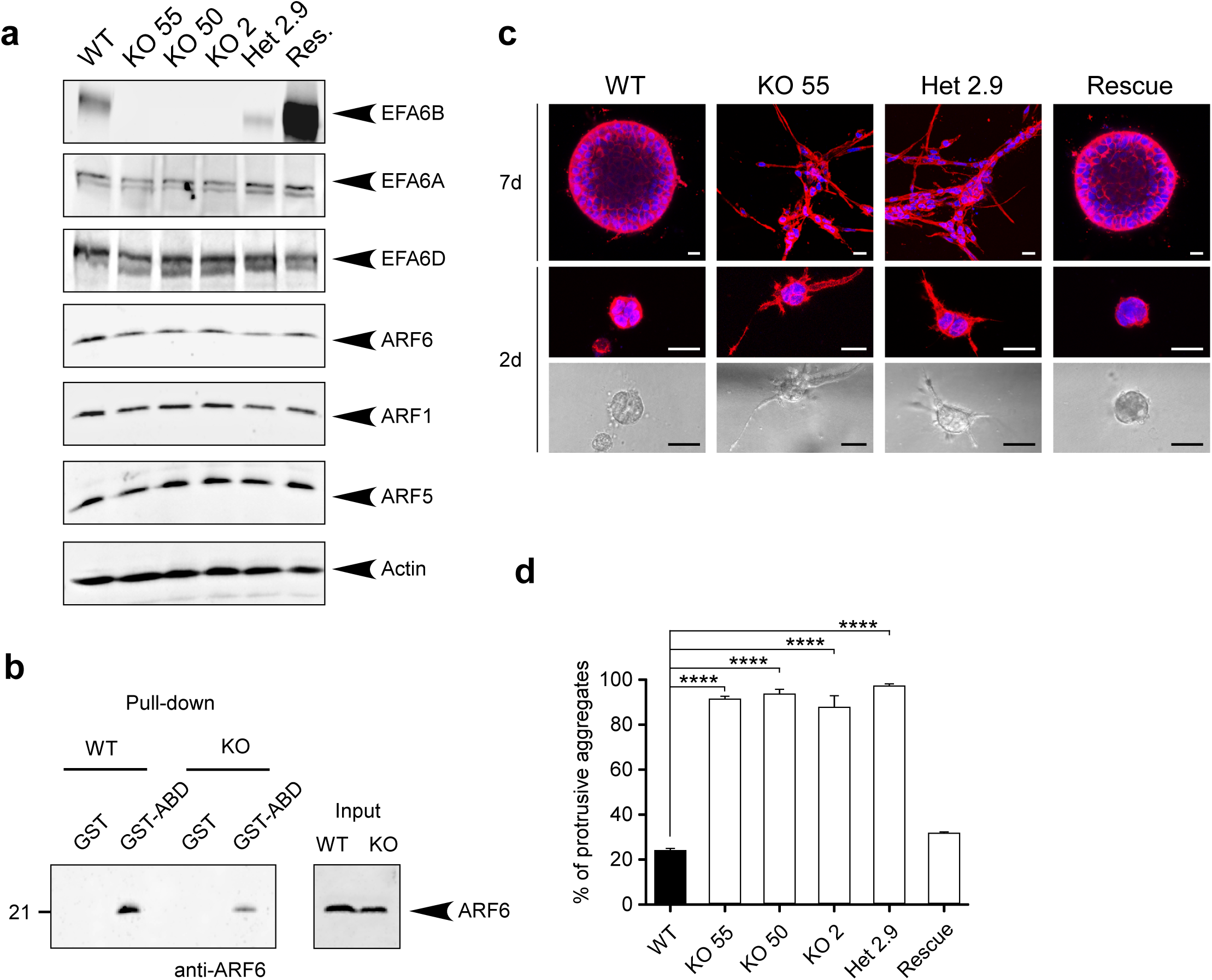
CRISPR/Cas9-mediated knock-out of the EFA6B encoding gene *PSD4* in MCF10A cells induces collective invasion in collagen I. **a**) The MCF10A WT, the homozygous EFA6B KO55, EFA6B KO50, EFA6B KO2, the heterozygous EFA6B KO2.9 and the EFA6B KO55 over-expressing EFA6B-vsvg cells (Rescue) were solubilized and the expression of the indicated proteins were analyzed by immunoblot. Actin served as a loading control. **b**) Lysates of MCF10A WT and EFA6B KO55 cells were reacted with GST or GST-ABD (ARF6GTP-binding domain of ARHGAP10) prebound to glutathione-sepharose beads. The whole lysates and bound proteins were analyzed by immunoblotting with an anti-ARF6 antibody. **c**) Representative images of the MCF10A WT, the homozygous EFA6B KO55, the heterozygous EFA6B KO2.9 and the EFA6B KO55 over-expressing EFA6B-vsvg placed in collagen for 7 days (upper panels) or 2 days (middle and bottom panels). The cells were processed for immunofluorescence to label the endogenous F-actin (red) and the nuclei (blue). The bottom panels are brightfield phase contrast images of the corresponding immunofluorescence images shown in the middle panels. Scale bars 20 μm. **d**) Quantification of the percentage of cell aggregates with invasive protrusions of the indicated MCF10A cell lines grown in collagen for 2 days. n=6, average ±SEM, Student’s *t*-test *p* values were ****, *p < 0.0001*.

To assess the hallmark property of epithelial cells to polarize and self-organize in acini, the clones were placed in collagen I gels. Control cells formed round aggregates typical of normal epithelial cells. In contrast, all the KO clones outgrew branched structures reminiscent of collective invasion (**Fig.1c**) Quantification of cell aggregates displaying at least one branched structure showed that KO clones developed 4 times more membrane and cellular protrusions than WT cells (**Fig.1d**), which was accompanied by neither increased cell proliferation nor migratory properties **(Supplementary Fig.1).** Re-expression of wild-type EFA6B by lentiviral infection was sufficient to recover the formation of normal round aggregates (**Fig.1a,c,d**) indicating that the invasive phenotype was specifically a consequence of the loss of EFA6B. The heterozygous (Het2.9) clone organized identical branched structures to the same extent as the homozygous KO clones (**Fig.1c,d**). Because exogenous expression of EFA6B in KO55 cells rescued the invasive phenotype, we propose that the dominant loss-of-function of EFA6B in the heterozygous KO cells is likely due to haplo-insufficiency.

### Loss of EFA6B stimulates invasive branching in luminal and basal mammary populations

We next asked whether the EFA6B KO has a similar impact on both luminal and basal mammary epithelial cells. We used the HMLE human epithelial population as it contains luminal progenitors and both mature luminal and basal epithelial cells. The markers EpCAM (Epithelial Cell Adhesion Molecule) and CD49f (ITGα6) are commonly used to assess mammary cell differentiation^9^. We sorted luminal progenitors (EpCAM+/CD49f+), mature luminal (EpCAM+/CD49f-) and mature basal (EpCAM-/CD49f+) cells (**Fig.2a**) and immediately performed CRISPR/Cas9-mediated invalidation of *PSD4* in each separate population. We obtained one homozygous (KO3) and one heterozygous (Het.25) KO clones from the luminal progenitor population and one homozygous clone (KO1) from the mature basal population. No clone was obtained from the mature luminal population. EFA6B protein expression was undetectable in the homozygous KO clones, while the heterozygous clone expressed half of its corresponding WT clone (**Fig.2b**). The expression of EFA6 paralogs, EFA6A and EFA6D, was unaffected. A significant reduction of ARF6 expression was observed in the clones isolated from the luminal progenitor population but not from the basal population. However, neither ARF1 nor ARF5 levels were altered. In 3D-collagen culture, HMLE WT clones formed cohesive rounded aggregates while the EFA6B homozygous and heterozygous KO clones displayed invasive cellular protrusions (**Fig.2c**). Although less branched compared to MCF10A, the HMLE KO clones formed invasive aggregates (**Fig.2d**). In conclusion, EFA6B is a general negative regulator of the invasive properties of epithelial mammary cells from both luminal and basal origins and loss of one *PSD4* allele results in haplo-insufficiency.

**Fig.2:**
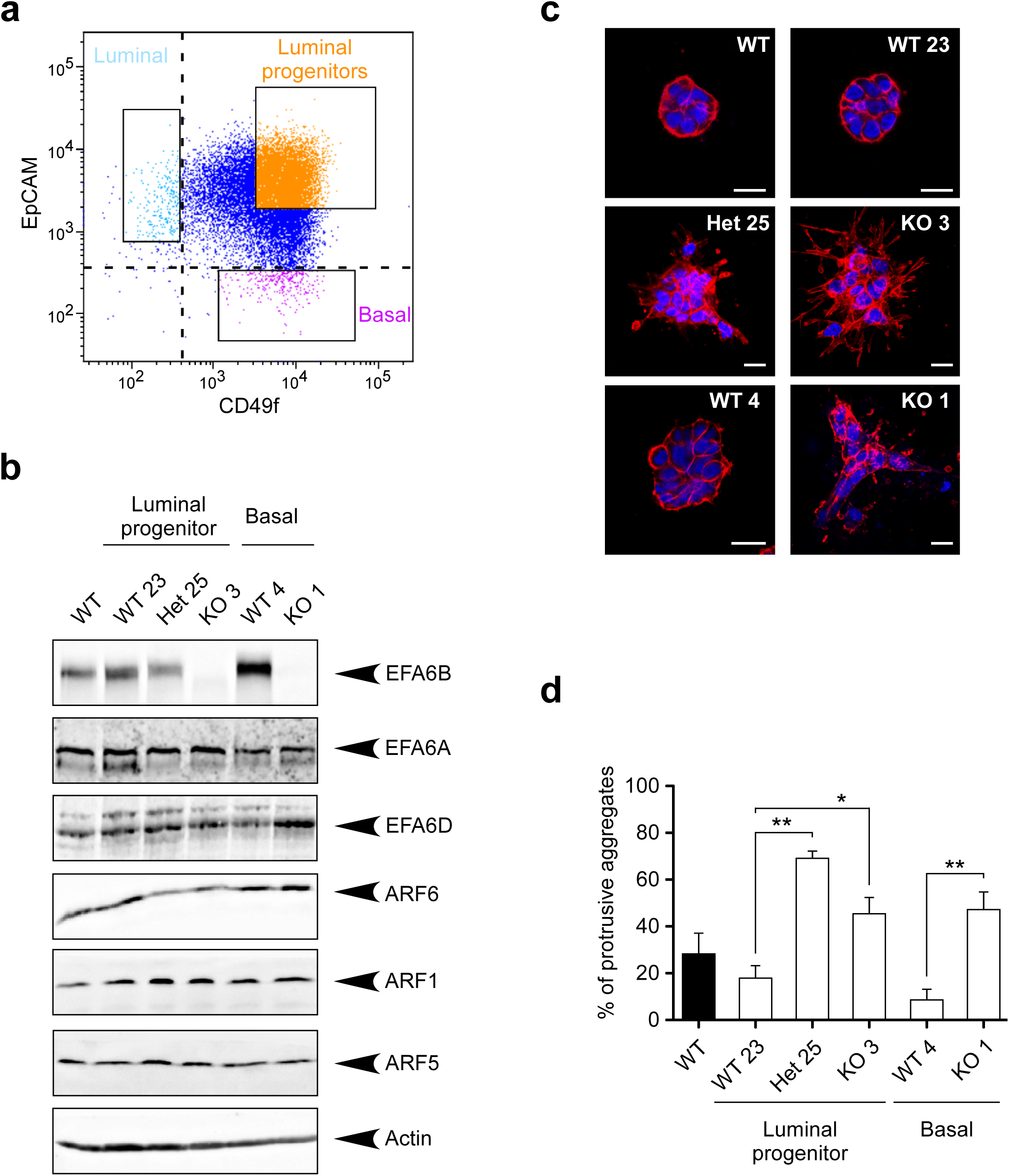
CRISPR/Cas9-mediated knock-out of the EFA6B encoding gene *PSD4* in HMLE luminal and basal populations induces collective invasion in collagen I. **a**) The cell surface marker EpCAM and CD49f were used to sort three epithelial cell populations including the luminal, luminal progenitors and mature basal cells. These cells were immediately processed for CRISPR/Cas9-mediated *PSD4* knock-out. **b**) The HMLE WT population, the luminal progenitor clone WT23, heterozygous EFA6B KO25, homozygous EFA6B KO3 and the mature basal clone WT4, homozygous EFA6B KO1 cells were solubilized and the expression of the indicated proteins analyzed by immunoblot. Actin served as a loading control. **c**) Representative images of the indicated cells grown 5 days in collagen I and stained for F-actin (red) and the nuclei (blue). Scale bars 20 μm. **d**) Quantification of the percentage of cell aggregates with invasive protrusions of the indicated cell lines grown in collagen I for 5 days. n=5, average ±SEM, Student’s *t*-test *p* values were * *p < 0.05*; **, *p < 0.01*.

### EFA6B knock-out stimulates invasion through the formation of MMP14-based degradative invadopodia

To determine whether EFA6B KO MCF10A cells acquired degradative properties, we seeded them on fluorescent gelatin matrix. In contrast to WT cells, the KO55 clone was capable of degrading the fluorescent gelatin seen as dark spots underneath the cells (**Fig.3a,b**). We also performed immunofluorescence analyses using the Col1-3/4C antibody that recognizes specifically the digested ends of collagen fibers^10^. The Col1-3/4C staining was virtually absent around the WT aggregates while a strong signal was visible along the invasive cellular protrusions extending from the KO55 aggregates (**Fig.3c,d**). Thus, the KO of EFA6B enabled MCF10A cells to proteolytically cleave collagen fibers organized as a 3D matrix. Searching for the protease responsible for the collagen degradation, we focused on MMP14 described as the main metalloprotease involved in mammary gland morphogenesis and BC metastasis^11–14^. We observed that down-regulation of MMP14 strongly inhibited collagen invasion by the KO cells (**Fig.3e,f**). Because the degradation of the fluorescent gelatin appeared as circular spots and because WT and KO55 cells globally expressed similar levels of MMP14 (**Fig.3g**), we searched for the formation of invadopodia enriched in MMP14. First, in EFA6B KO cells, staining for cortactin, a well-known marker for invadopodia, revealed the presence of large structures co-stained for F-actin (**Fig.3h and Supplementary Fig.4**). In WT cells, the cortactin positive spots, which were not as enriched in F-actin, were smaller. In addition, the cortactin positive aggregates observed in EFA6B KO cells co-localized with the black spots of degraded fluorescent gelatin **(Supplementary Fig.1)**. Quantification of the percentage of cells with invadopodia showed a 2-fold increase in KO cells when compared to WT cells (**Fig.3i and Supplementary Fig.4**). We then looked for MMP14 localization in invadopodia. Cells were transiently transfected with a MMP14-mCherry encoding plasmid and stained for cortactin. KO55 cells, but not WT cells, expressing MMP14-mCherry presented extensive co-localization with cortactin (**Fig.3j**). These results demonstrate that the loss of EFA6B leads to the formation of degradative MMP14-enriched invadopodia responsible for invasion within collagen I.

**Fig.3:**
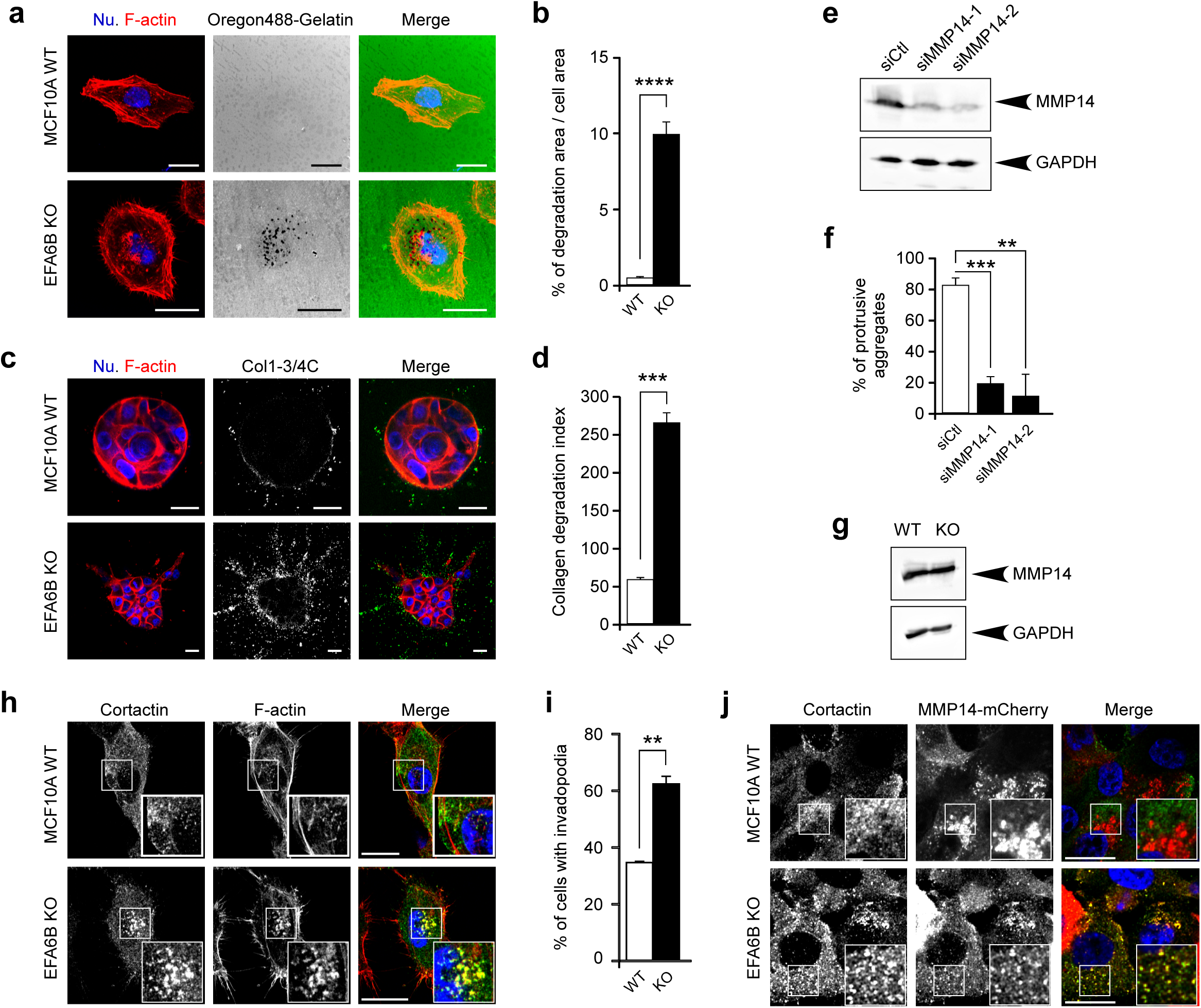
EFA6B knock-out stimulates matrices degradation and invasion in a MMP14-dependent manner. **a**) Representative images of MCF10A WT and EFA6B KO55 cells placed on Oregon488-gelatin (green) coated coverslips and stained for F-actin (red) and nuclei (blue). Areas devoid of fluorescent signal indicate degradation of the fluorescent gelatin. Scale bars 20 μm. **b**) Quantification of the gelatin degradation. Values are mean percentage of degradation area per cell area ± SEM. 90 cells were analyzed for each cell population in three independent experiments. ****, *p < 0.0001*. **c**) Representative images of MCF10A WT and EFA6B KO55 cells grown in collagen I for 3 days and stained for cleaved collagen I with the Col1-^3/4^C antibody (white in middle panel and green in left merge panel), for F-actin (red) and nuclei (blue). Scale bars 20 μm. **d**) Quantification of the collagen degradation. Values are mean degradation index ± SEM. 90 cells were analyzed for each cell population in three independent experiments. ***, *p < 0.001*. **e**) MCF10A WT and EFA6B KO55 cells were transfected with siRNA control or directed against MMP14. 48h post-transfection the expression of MMP14 was analyzed by immunoblot. GAPDH served as a loading control. **f**) Quantification of the percentage of cell aggregates with invasive protrusions of the indicated cell lines grown in collagen I for 2 days. n=4, ****, *p < 0.0001*. **g**) Expression of MMP14 analyzed by immunoblot in MCF10A WT and EFA6B KO55 cells. GAPDH served as a loading control. **h**) Representative images of the indicated cells grown 2 days on collagen I-coated coverslips stained for cortactin (green), F-actin (red) and nuclei (blue). The large inset is a 2X zoom-in image of the indicated area. Scale bars 20 μm. **i**) Quantification of the percentage of cells displaying invadopodia. **j**) Representative images of the indicated cells grown 2 days on collagen I-coated coverslips stained for cortactin (green), MMP14-mCherry (red) and nuclei (blue). The large inset is a 2X zoom-in image of the indicated area. Scale bars 20 μm.

### EFA6B knock-out promotes a change in the ITG repertoire and stimulates the formation of ITGβ1 - based degradative invadopodia

Invasion and formation of invadopodia relies on the ITG-mediated interaction with the ECM^5^. To assess the changes promoting invasion imposed upon loss of EFA6B, we compared the gene expression profile of *PSD4* KO MCF10A cells (N=5, including KO55 (N=3) and KO2.9 cell (N=2)) to that of *PSD4* WT MCF10A cells (N=3). We identified 296 genes differentially expressed, including 173 overexpressed and 123 underexpressed in the KO cells (**Supplementary Table 1**). Of note, the matrisome and its receptor machineries were the main affected group of genes with up to 12% (36 genes) of the total 296 altered genes: affected matrisome genes encoded structural glycoproteins, ECM-associated proteins, regulators and secreted factors, as well as ECM receptor machinery molecules. Indeed, many gene ontologies associated to this gene list were related to ECM, cell-cell signaling, cell-cell adhesion, cell-adhesion (to substratum) and EMT **(Supplementary Table 2)**. GSEA of a “Matrisome+receptors” gene set (including the KEGG gene sets Focal Adhesion (“FA”), ECM-receptor interaction (‘ECM”) and the human matrisome database confirmed such over-representation of genes **(Supplementary Fig.2).** Thus, in response to *PSD4* mutation the cells have modified their molecular extracellular environment and the expression of corresponding receptors.

We then explored the protein expression level of the major ITGs that regulate mammary morphogenesis. We found that KO55 cells have a reduction of subunits α6 (50%) and β4 (36%) expression, while subunits α2, α3 and β1 were unchanged (**Fig.4a-c**). To determine which ITGs were required for EFA6B KO-mediated invasion, we quantified the number of invasive aggregates grown in gels containing control or ITG-blocking antibodies. Antibodies against α6, β4 or α3 had no or little effect. In contrast, antibodies against α2 and β1 strongly blocked cell invasion suggesting that the ITGα2β1 is required for EFA6B KO cells invasion (**Fig.4d**). Since, the ITGβ1 has been involved in invadopodia regulation and MMP14 enrichment^15^, we assessed whether β1 was concentrated at invadopodia. We found a large co-localization of β1 with MMP14-mCherry in KO55 but not in MCF10A WT cells (**Fig.4e**). Thus, the absence of EFA6B induces a modification of the matrisome and ITG repertoire along with the formation of degradative ITGβ1-based invadopodia, all of which contribute to the collective invasion of the EFA6B KO cells.

**Fig.4:**
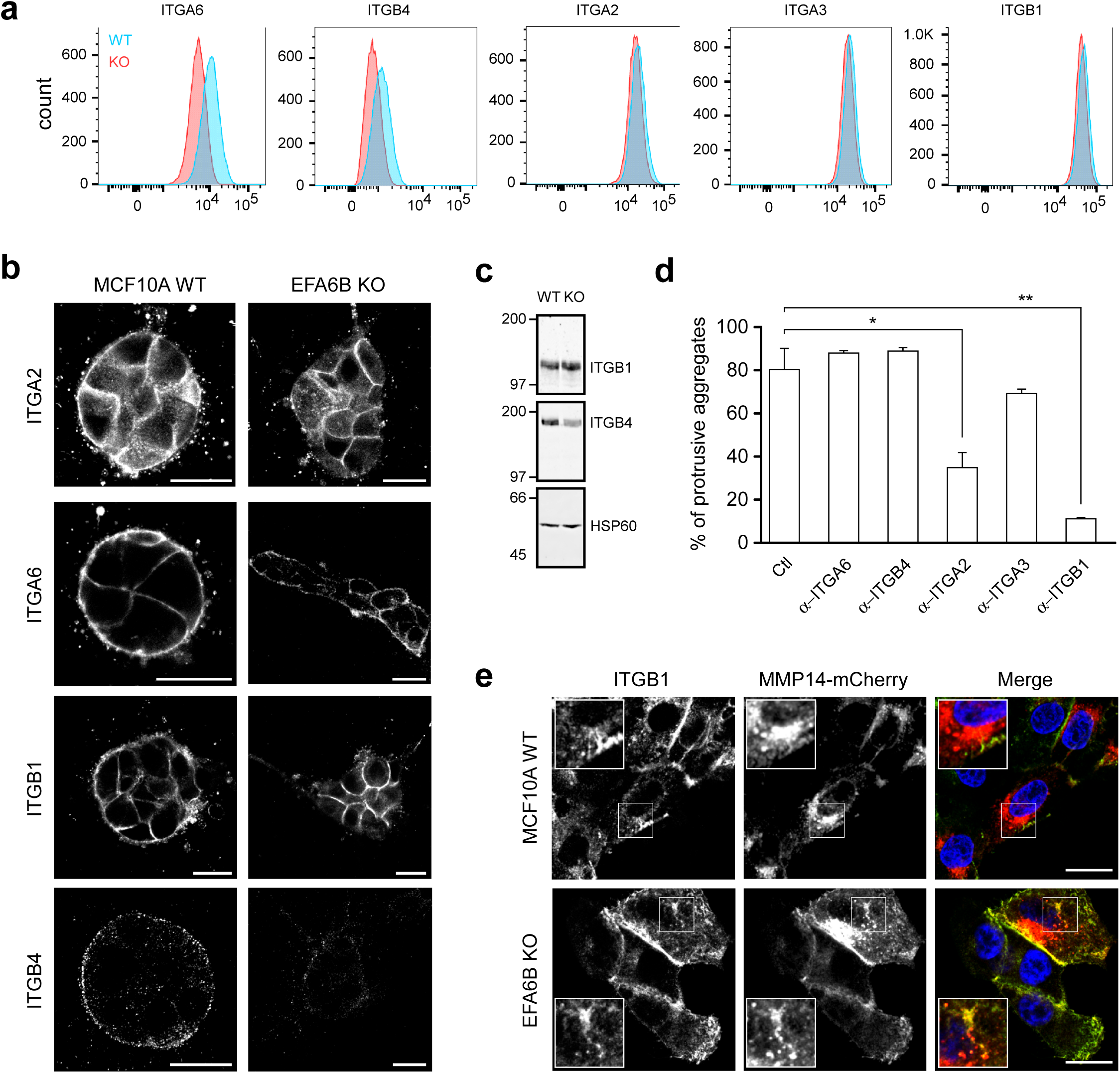
EFA6B knock-out promotes a change in the ITG repertoire and stimulates the formation of ITGβ1-based invadopodia. **a**) Cell surface expression of ITG molecules in MCF10A WT and EFA6B KO55 cells analyzed by FACS. **b**) Representative images of the indicated spheroids grown 2 days in collagen I stained for the indicated ITG. Scale bars 20 μm. **c**) Expression of ITGβ1 and ITGβ4 analyzed by immunoblot in MCF10A WT and EFA6B KO55 cells. HSP60 served as a loading control. **d**) Quantification of MCF10A WT and EFA6B-KO55 cell aggregates with invasive protrusions incubated in the presence of the control pre-immune serum (Ctl) or the indicated anti-ITG antibodies for 2 days. Values are percentages of total cell aggregates ± SEM. 300 cell aggregates were analyzed for each cell population in three independent experiments. *, *p < 0.05*; **, *p < 0.01*. **e**) Representative images of the indicated cells grown 2 days on collagen I-coated coverslips stained for ITGβ1 (green), MMP14-mCherry (red) and nuclei (blue). The large inset is a 2X zoom-in image of the indicated area. Scale bars 20 μm.

### EFA6B knock-out induces the expression of EMT transcription factors that promote collective invasion of MCF10A and HMLE cells in collagen I

EMT is a major molecular program promoting collective invasion believed to provide cells with migratory and degradative advantages^5,16^. We observed that the MCF10A EFA6B KO cells had reduced levels of E-cadherin together with an increased expression of N-cadherin (**Fig.5a and Supplementary Fig.3**). Knowing that EFA6B is a tight junction (TJ) regulator, we also looked at TJ molecules and found that CLDN1 and CLDN3 were strongly reduced in all clones and occludin decreased in 3 out of 4 clones (**Fig.5a**). We assessed the cell-to-cell affinity by using the hanging-drop assay. MCF10A WT cells formed compact monolayers while KO55 cells formed lacy monolayers indicative of loose cell-cell contacts (**Supplementary Fig.3**). In addition, our transcriptomic analysis of the KO55 and Het2.9 clones showed a significant alteration of the expression of the Gene Ontology “cell-cell adhesion” signature (**Supplementary Fig.2 and Supplementary Table 2**). These results support a mode of collective invasion whereby reduced cell-cell adhesion facilitates pro-migratory cell movements while maintaining cell-cell contacts.

**Fig.5:**
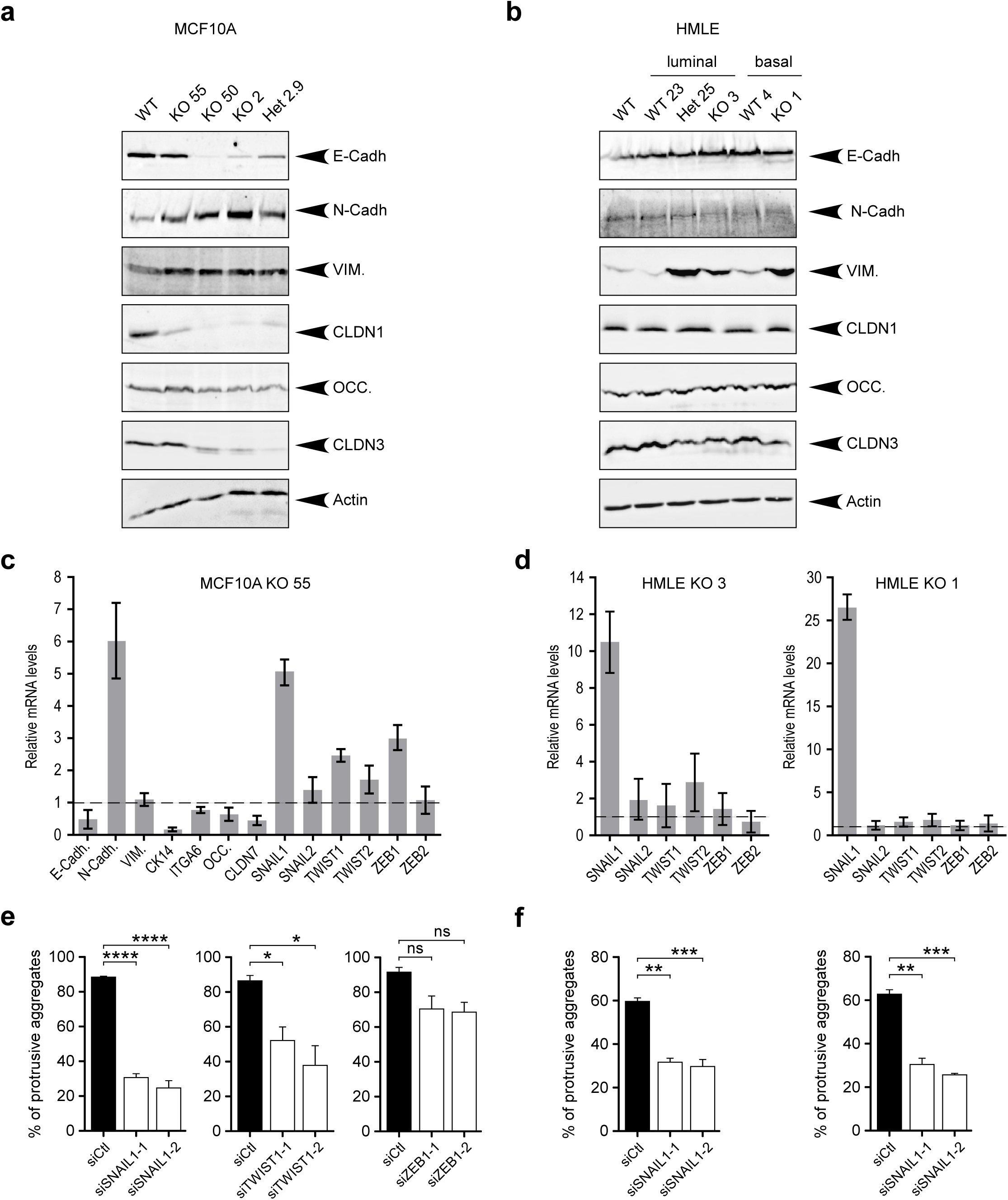
EFA6B knock-out induces the expression of EMT transcription factors that promote collective invasion of MCF10A and HMLE WT cells in collagen I. **a**) The MCF10A WT, the homozygous EFA6B KO55, KO50, KO2 and the heterozygous EFA6B KO2.9 cells were solubilized and the expression of the indicated proteins analyzed by immunoblot. Actin served as a loading control. **b**) Expression of EMT-associated genes by qPCR analysis in EFA6B KO55 cells normalized to MCF10A WT. n=3, average ±SEM. **c**) The HMLE WT population, the luminal progenitor clone WT23, heterozygous KO25, homozygous KO3 and the mature basal clone WT4, homozygous KO1 cells were solubilized and the expression of the indicated proteins analyzed by immunoblot. Actin served as a loading control. **d**) Expression of EMT-TF genes by qPCR analysis in EFA6B KO3 and KO1 cells normalized to their respective HMLE control cells WT23 and WT4. n=6, average ±SEM. **e**) Quantification of the percentage of aggregates with invasive protrusions of MCF10A KO55 cells grown in collagen I for 2 days after transfection with the indicated siRNAs. n=3, average ±SEM, Student’s *t*-test *p* values were * *p < 0.05*; ****, *p < 0.0001*, non significant (ns). **f**) Quantification of the percentage of aggregates with invasive protrusions of HMLE KO3 (left) and KO1 (right) cells grown in collagen I for 2 days after transfection with the SNAIL1 targeted siRNAs. n=3, average ±SEM, Student’s *t*-test *p* values were **, *p < 0.01*; ***, *p < 0.001*.

Further analysis by RT-qPCR confirmed the E/N-cadherin switch, the decrease of TJ markers and, of *CK14* whose down-regulation was recently shown to mark an advanced mesenchymal state in melanoma^17^, and the decrease of the mammary differentiation marker *CD49f* (**Fig.5b**). We found an overall decrease of EpCAM and CD49f (66.9±7.2% and 41.4±9.6%, respectively), together with the emergence of a new EpCAM^-/low^ and CD49f^low^ population (11.7% of total cells, red gate), suggestive of a loss of epithelial identity of the KO55 cells (**Supplementary Fig.3**).

We similarly analyzed the EMT status of the HMLE EFA6B KO clones. Although we did not notice a change in E- or N-cadherin expression, we found a strong increase of vimentin and a slight but consistent decrease of CLDN3 expression (**Fig.5c**). Because of the well-documented variable outcome of EMT^16^, a more recent definition of EMT relies on the turn-on of EMT-activating transcription factors (EMT-TFs)^18^. We found a significant increase of *SNAIL1, TWIST1* and *ZEB1*, and to a lower extent of *SNAIL2* and *TWIST2* expression in the KO55 cells (**Fig.5b**). In HMLE, *SNAIL1* was the major EMT-TF commonly up-regulated in all three KO clones (**Fig.5d and Supplementary Fig.3**). Hence, depending on the cell lines, the elicited EMT-TFs and the EMT program triggered by EFA6B KO are variable, yet altogether our results indicate that the loss of EFA6B engages epithelial cells into EMT.

Activation of EMT-TFs was shown to favor dissemination by increasing invasion^5,16^. We found that siRNA-mediated down-regulation of *SNAIL1* in KO55 cells blocked invasion, while that of *TWIST1* only reduced it and that of *ZEB1* had no effect (**Fig.5e and Supplementary Fig.3**). Similarly, depletion of *SNAIL1* in both HMLE KO clones blocked invasion (**Fig.5f and Supplementary Fig.3**), indicating that SNAIL1 is the major EMT-TF required for collagen invasion of EFA6B KO cells.

EMT has been found to be associated with a gain in stemness properties ^19,20^. Analyses of the MCF10A clones failed to detect the formation of any mammosphere and showed no gain in the mammary stem cell characteristics **(Supplementary Fig.3).**

We concluded that the knock-out of EFA6B in mammary cells induces the expression of EMT-TFs triggering an EMT program that promotes dedifferentiation and facilitates collective invasion.

### EFA6B knock-out stimulates cellular contractility and invasion through CDC42

ECM remodeling and cellular contractility are important regulators of the formation of invadosomes^21,22^. We first studied the impact of EFA6B KO on cell contractility by using a collagen gel contraction assay. KO55 cells were at least twice more contractile than their WT counterparts (**Fig.6a**). Cell contractility is dependent upon the activation of the myosin II through the phosphorylation of the myosin light chain (MLC)^23^. KO55 cells presented higher levels of pMLC than control cells (**Fig.6b**). We then looked at the organization of the fibrillary collagen surrounding the cell aggregates by reflectance (**Fig.6c**). We could not distinguish any particular orientation of the collagen fibers, which were randomly distributed around the WT cell aggregates. In contrast, reflectance signals showed a radial distribution of aligned collagen fibers in KO55 invasive cells. In addition, the collagen fibers were in perfect alignment with membrane filopodial structures extended from the cells (**Fig.6c**). This observation indicated that EFA6B KO cells have the capacity to remodel the collagen fibers into tracks to facilitate their invasion. Small G proteins of the RHO family are known to control cell contractility ^24^. We found that the downregulation of CDC42, but not RHOA or RAC1, altered the contractility of the KO55 and KO2 cells (**Fig.6a,d and Supplementary Fig.4**). Next, we observed that the down-regulation of CDC42 was the most effective at blocking invasion, while RHOA and RAC1 had no effect **(Fig.6d,e)**. It also hampered the formation of invadopodia and the extension of filopodia (**Fig.6f,g and Supplementary Fig.4**). In agreement with the functional assays, we found an upregulation of activated CDC42 (2.8±0.8), but not of RHOA (1.1±0.1) nor RAC1 (0.9±0.10) (**Fig.6h**). Overall, these observations demonstrated that EFA6B KO leads to the activation of CDC42, which regulates cell contractility, formation of filopodia and invadopodia, collagen remodeling and collective invasion in 3D-collagen.

**Fig.6:**
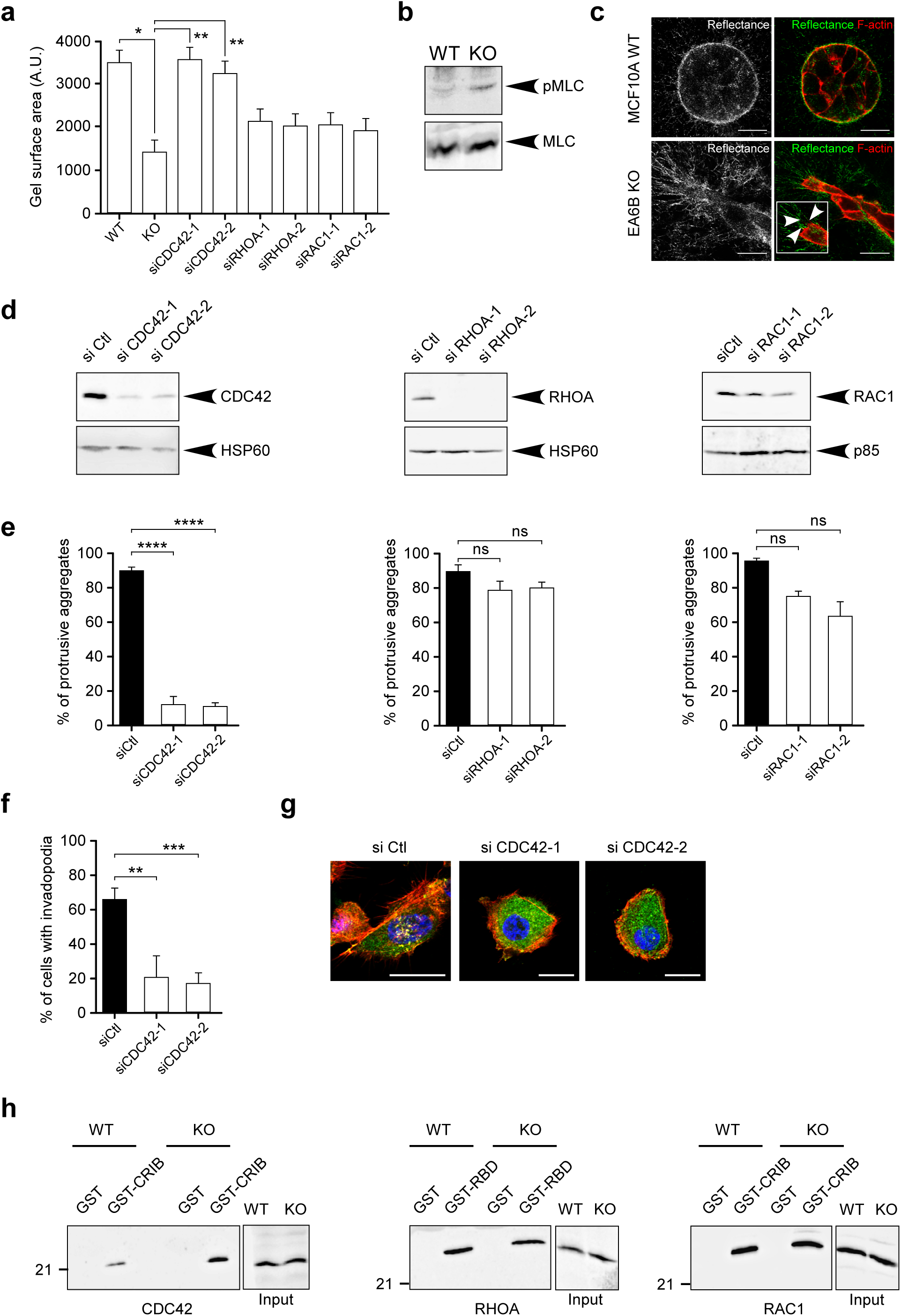
EFA6B knock-out stimulates cellular contractility and invasion through CDC42. **a**) Quantification of the contractility of MCF10A WT (WT) and EFA6B KO55 (KO) cells transfected with a control siRNA, and EFA6B KO55 cells transfected with the indicated siRNA evaluated by a collagen gel contraction assay. Values are mean surface area of the collagen gel ± SEM. n*=3*, Student’s *t*-test *p* values were * *p < 0.05*; ** *p < 0.01*. **b**) Expression of pMLC and total MLC analyzed by immunoblot in MCF10A WT and EFA6B KO55 cells. **c**) Representative images of the MCF10A WT and EFA6B KO55 spheroids embedded 2 days in collagen I stained for F-actin (red). The organization of the collagen I fibers surrounding the cell aggregates was imaged by confocal reflectance microscopy (green). The large inset is a 2X zoom- in image of the leader cell. Arrowheads point to thin membrane extensions co-localized with collagen fibers. Scale bars 20 μm. **d**) Two days post-transfection of EFA6B KO55 cells with the indicated siRNAs, the expression of the corresponding protein was analyzed by immunoblot. HSP60 and p85 served as loading controls. **e**) Quantification of the percentage of aggregates with invasive protrusions of EFA6B KO55 cells grown in collagen I for 2 days after transfection with the indicated siRNAs. n=3, average ±SEM, Student’s *t*-test *p* values were **** *p < 0.0001*; ns: non-significant. **f**) Quantification of the percentage of invadopodia in EFA6B KO55 cells grown in collagen I for 2 days after transfection with CDC42 targeted siRNAs. n=3, average ±SEM, Student’s *t*-test *p* values were **** *p < 0.0001*. **g**) Representative images of EFA6B KO55 cells transfected with the indicated siRNAs and stained for cortactin (green), F-actin (red) and nuclei (blue). Scale bars 20 μm. **h**) Lysates of MCF10A WT and EFA6B KO55 cells were reacted with GST, GST-CRIB (CDC42GTP- and RAC1GTP-intercating domain of PAK) or GST-RBD (RHOAGTP-binding domain of rhotekin) prebound to glutathione-sepharose beads. The whole lysates and bound proteins were analyzed by immunoblot with the indicated antibodies.

### EFA6B knock-out stimulates cellular contractility and invasion through CDC42-MRCK-pMLC and CDC42-NWASP-ARP2/3 pathways

CDC42 was shown to regulate contractility through the recruitment of kinases that phosphorylate MLC^25,26^, as well as to control invadopodia formation by activating the ARP2/3 complex through N-WASP^27^. Downstream of CDC42, the myotonic dystrophy-related CDC42-binding kinases (MRCK) phosphorylate directly or indirectly MLC^28^. SiRNA experiments showed that the combined down-regulation of MRCKα/β in KO55 cells reduced both MLC phosphorylation and cell invasion in collagen I (**Fig.7a**). RHOA activates the RHO*-*associated protein kinases ROCK1 and ROCK2 to phosphorylate MLC^29,30^. Knock-down of both ROCK1/2 had no effect on either phosphorylation of MLC or invasion (**Fig.7b**). Thus, in KO55 cells, activated CDC42 controls phosphorylation of MLC and invasion in an MRCKα/β-dependent manner. Further, siRNA down-regulation of N-WASP and ARP3 inhibited invasion (**Fig.7c,d and Supplementary Fig.4**). Altogether, these results unraveled a new pathway regulated by EFA6B, which controls the activation of CDC42 and two of its effector functions: contractility through MRCKα/β phosphorylation of MLC and invasion through N-WASP and ARP2/3.

**Fig.7:**
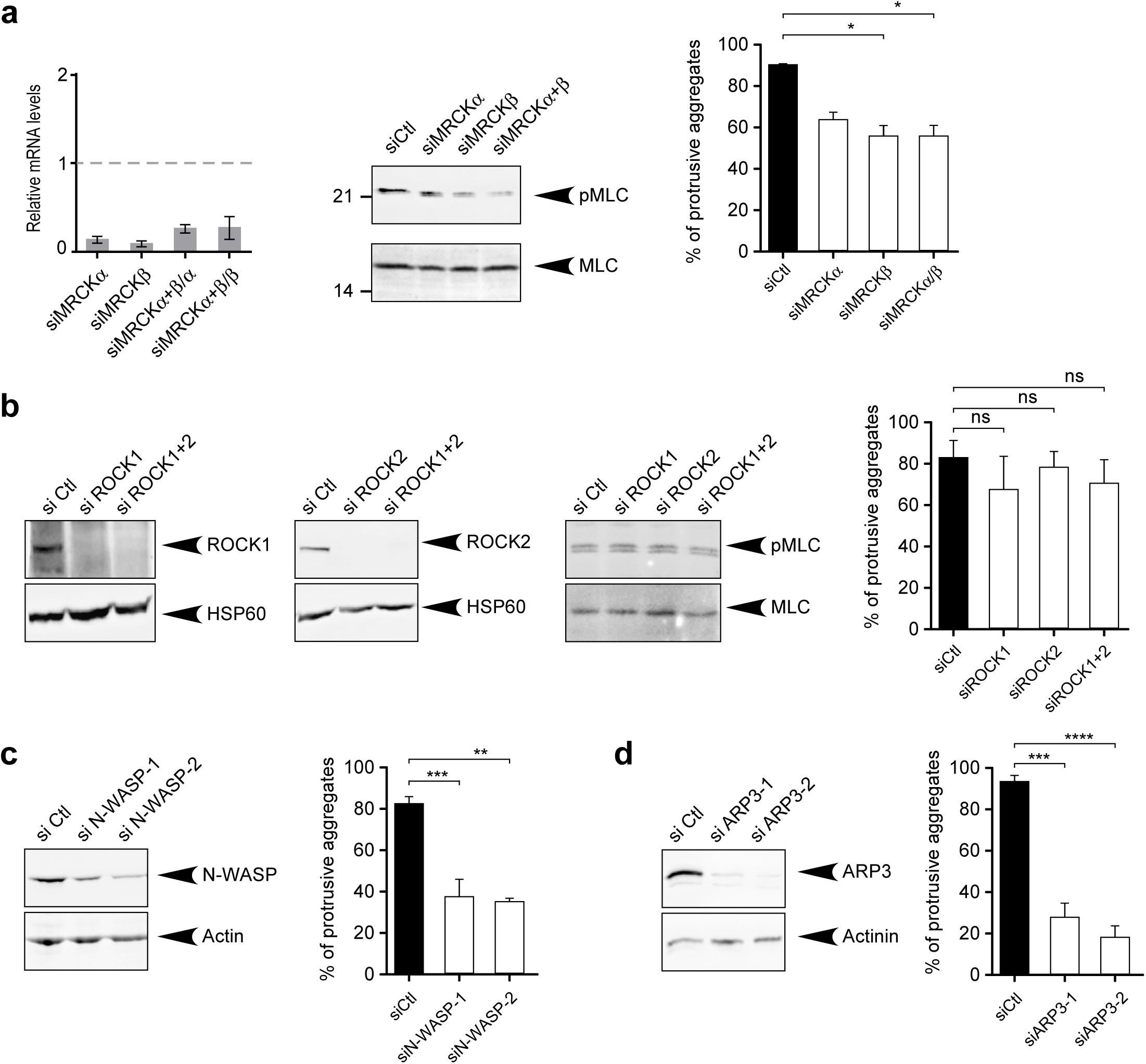
EFA6B knock-out stimulates cellular contractility and invasion through two CDC42-dependent signaling pathways: CDC42-MRCK-pMLC and CDC42-N-WASP-ARP2/3. **a**) Left: expression of MRCKα and MRCKβ genes in EFA6B KO55 cells analyzed by qPCR 2 days post-transfection with the indicated siRNAs. siMRCKα+β/α and siMRCKα+β/β indicate the gene expression level of MRCKα or MRCKβ after transfection with siRNAs against both MRCKs. Middle: expression of pMLC and total MLC analyzed by immunoblot two days post-transfection of EFA6B KO55 cells with the indicated siRNAs. Right: quantification of the percentage of aggregates with invasive protrusions of EFA6B KO55 cells grown in collagen I for 2 days post-transfection with the indicated siRNAs. n=3, average ±SEM, Student’s *t*-test *p* values were * *p < 0.05*. **b**) Expression of the indicated proteins analyzed by immunoblot two days post-transfection of EFA6B KO55 cells with the indicated siRNAs. HSP60 served as a loading control. Right panel: quantification of the percentage of aggregates with invasive protrusions of EFA6B KO55 cells grown in collagen I for 2 days after transfection with the indicated siRNAs. n=3, average ±SEM, Student’s *t*-test *p* values were non-significant (ns). **c**) Left: expression of N-WASP analyzed by immunoblot two days post-transfection of EFA6B KO55 cells with N-WASP targeted siRNAs. Actin served as a loading control. Right: quantification of the percentage of aggregates with invasive protrusions of EFA6B KO55 cells grown in collagen I for 2 days after transfection with N-WASP-directed siRNAs. n=3, average ±SEM, Student’s *t*-test *p* values were ** *p < 0.01*; ***, *p < 0.001*). **d**) Left: expression of ARP3 analyzed by immunoblot two days post-transfection of EFA6B KO55 cells with ARP3 targeted siRNAs. Actinin served as a loading control. Right: quantification of the percentage of aggregates with invasive protrusions of EFA6B KO55 cells grown in collagen I for 2 days after transfection with ARP3 targeted siRNAs. n=3, average ±SEM. Student’s *t*-test *p* values were *** *p < 0.001*; ****, *p < 0.0001*.

### EFA6B expression is down regulated in human clinical BC samples endowed with invasive properties

Our results demonstrate that the reduction of EFA6B expression in normal mammary cells is sufficient to promote the acquisition of invasive properties. Within transformed cells the loss of EFA6B could provide a pro-invasive advantage favoring the transition from *in-situ* to invasive lesion. To assess the impact of EFA6B down-regulation in BC progression, we compared the *PSD4* mRNA expression levels in unpaired clinical samples of ductal carcinoma *in situ* (DCIS) and invasive ductal carcinoma (IDC)^31,32^. We found a significant reduction of *PSD4* expression, but not of the other family members, in IDC compared to DCIS (**Fig.8a**). We then analyzed separately tumor epithelium and adjacent stroma prepared by laser-capture microdissection for each of the 22 unpaired DCIS (N=11) and IDC (N=11)^31^. A significant reduction of *PSD4* levels was observed only in the epithelial compartment (**Fig.8b**), which corroborates our results showing that the loss of EFA6B in cells of epithelial origin generates invasive mammary cells **(Fig.2)**. Interestingly, several GO associated with our 296-gene signature and related to ECM organization, cell-cell adhesion and EMT (**Fig.8c**), were also associated with the genes identified by Lee *et al.* as differentially expressed in IDC versus DCIS^31–33^ and related to disease progression. Thus, ontologies associated with *PSD4* knock-out in MCF10A cells correlate with those of the progression of DCIS to IDC in patients. We also compared the *PSD4* mRNA expression in metastatic samples versus the paired primary BC by taking advantage of a publicly available series of 29 matched metastasis/primary cancer pairs^34,35^. We found significant reduction of *PSD4* expression in the metastatic samples (**Fig.8d**), further corroborating the role of EFA6B loss in tumor invasion.

**Fig.8:**
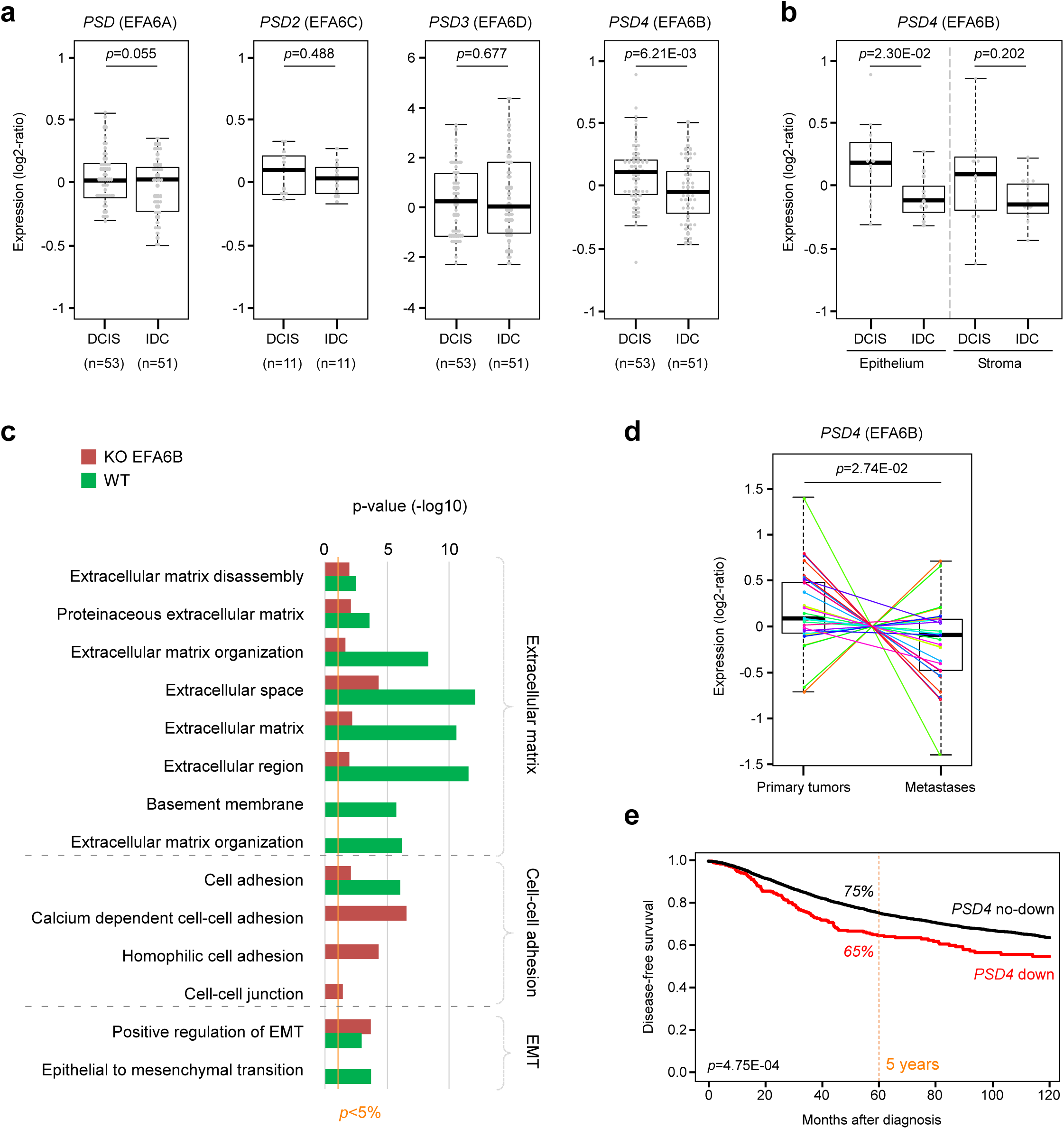
EFA6B/PSD4 expression is down-regulated in human clinical BC samples endowed with invasive properties. **a**) Expression of the four EFA6 isoforms in DCIS compared to IDC. The *p*-values are for the Student’s *t*-test. **b**) Expression of *PSD4* is selectively decreased in the epithelial compartment of DCIS (n=11). The *p*-values are for the Student’s *t*-test. **c**) Ontologies associated with *PSD4* knock-out in MCF10A cells. **d**) Expression of *PSD4* is decreased in the metastatic samples *versus* the paired primary BC samples (n=29). The corresponding primary BC–metastasis pairs are connected by thin colored lines. The *p*-value is for the paired Mann-Whitney test. **e**) Kaplan-Meier curves for DFS according to *PSD4* expression. The *p*-value is for the log-rank test.

Finally, to confirm and extend our previous results^7^ on a larger series, we searched for correlation between *PSD4* mRNA expression and the clinico-pathological features of our updated large publicly available series of 8,464 invasive primary BC (**Supplementary Table 3**). From this cohort, a total of 530 tumors showed a two-fold or greater down-regulation of *PSD4*, using normal breast tissue as the standard. *PSD4* down-regulation was associated (Fisher’s exact test; **Supplementary Table 4**) with higher pathological grade (p<0.001) and tumor size (p<0.001), ductal type (p=0.002), higher frequency of ER-negative (p<0.001) and ERBB2-negative (p<0.001) statutes, and shorter disease-free survival (DFS). Within the 6,156 non-stage IV patients with follow-up available, the 5-year DFS was 75% (95CI, 74-76%) for the whole population, and 65% (95CI, 59-70%) and 75% (95CI, 74-77%) in cases of down-regulation and no down-regulation respectively (p=4.75E-04, log-rank test; **Fig.8e**).

We conclude that loss of EFA6B endows epithelial mammary cells with molecular characteristics of human invasive ductal carcinoma.

## Discussion

### EMT

The invasive process occurring during embryonic development or cancer is believed to be associated with the activation of an EMT program. We observed that EFA6B KO cells generated from MCF10A and HMLE cells had an increased expression of EMT-TFs and displayed altered expression of classic EMT proteins. EMT is not a uniform program executed by a unique signaling pathway, and thus the differences between MCF10A and HMLE EFA6B KO cells likely arise from intrinsic properties to respond to EFA6B loss and carry on EMT. In addition, we observed that the basal and luminal progenitor HMLE KO clones were all invasive, and that regardless of their basal or luminal origin, they displayed similar EMT profiles. Thus, EFA6B depletion in both luminal and basal cells induces EMT and stimulates collective invasion. In agreement with our results, it was recently reported that the endocytic protein NUMB inhibits the acquisition of migratory and invasive properties by activating EFA6B/ARF6 in response to growth factors^36^.

### Mode of invasion

We found that our EFA6B KO cells assemble degradative ITGβ1-invadopodia enriched in MMP14. A large body of evidence obtained from *in vitro* and *in vivo* studies demonstrated that the MMP14 metalloprotease is instrumental in enabling normal mammary gland development and BC collective invasion^11–13,37^.

Interestingly, the fact that the EFA6B KO cells undergo EMT, display thin filopodia at the front of the leader cells, assemble MMP14 invadopodia and sustain invasive protrusions in collagen I is reminiscent of the mechanism of invasion by cancer cells.

Indeed, EMT program bestows carcinoma cells many of the attributes associated with invasive malignant cells^5,16^. Furthermore, collective invasion during mammary tubulogenesis occurs without actin-based protrusions extending into ECM^38^. Also, MMP14 is not expressed by the normal epithelial cells during mammary gland branching but is expressed by the invading BC cells^12^. Finally, normal cells would only invade collagen I transiently before producing their basement membrane that blocks invasion^6^. Thus, we conclude that the collective invasion of EFA6B KO normal mammary cells models cancer invasion rather than normal mammary branching.

### The CDC42 signaling pathway

The depletion of EFA6B led to a strong activation of CDC42 but not RAC1 nor RHOA, though largely involved in invasion and contractility^39^. *In vitro* and *in vivo* studies have shown important roles for CDC42 in regulating diverse cellular processes such as cell cycle progression and mitosis, polarity, survival, differentiation and stem cell function^40^. In the mammary gland, transgenic overexpression of CDC42 produced hyperbranched ductal trees with abnormal acini^41^, and in breast tumors CDC42 is often found hyperactivated or over-expressed^42,43^. However, activating mutations in CDC42 have not been found^44^, which suggests that its activating pathways must be altered. We propose that the loss of expression of EFA6B interferes with CDC42 regulation contributing to its hyperactivation in BC. From *in vitro* studies, CDC42 is a known pro-invasive protein that can induce the formation of filopodia, invadopodia and can regulate cellular contractility^40^. Consistent with this, we found that knock-down of CDC42 abrogates contractility and invasion of the EFA6B KO cells. CDC42 promoted increased contractility through MLC phosphorylation via the MRCK proteins. In contrast, the main RHOA/ROCK contractility pathway did not appear to play any role. Contractility is reflected by the formation of linearized bundles of collagen extending from filopodia of invasive EFA6B KO cells. These collagen bundles were shown to form tracts that facilitate tumor cells migration^21,45^. Cell contractility stiffens the matrix, and thus stimulates the formation of invodopodia and secretion of MMPs^46^. Interestingly, ECM rigidity and cellular contractility were shown to regulate the formation of invadopodia^15,47^, advocating that in EFA6B KO cells CDC42 activation facilitates the formation of invadopodia, at least in part, by increasing cell contractility. The increased contractility of EFA6B KO cells may also be the consequence of an alteration of the matrisome that might have strengthen the ECM rigidity, for instance through the observed up-regulation of the LOX protein^21^ (**Supplementary Table 1**).

Further, CDC42 plays a direct role in invadopodia formation^40,48^ and maturation by maintaining the activation of N-WASP-ARP2/3 complex, which in turn nucleates actin polymerization^49^. In addition, CDC42 promotes the recruitment of MMP14 to invadopodia^50^. Since the invasive nature of the EFA6B KO cells is dependent on N-WASP, ARP2/3 and MMP14, we conclude that CDC42 is the major effector of this behavior.

### EFA6B haplo-insufficiency and BC

Heterozygous KO clones were as invasive as the homozygous ones, emphasizing the importance of the levels of expression of EAF6B in repressing the invasive potential of normal mammary cells. This observation is in agreement with many of the results we have previously obtained by studying EFA6A and EFA6B in different cellular models indicating that their expression levels is a critical determinant of their physio-pathological functions^7,51–53^. Our results show that *PSD4/EFA6B* behaves as a haplo-insufficient gene with respect to the acquisition of an invasive phenotype. Since in BC patients we found a correlation between the severity of the clinico-pathological data and the gradual loss of expression of *EFA6B* messenger^7^, we propose that *PSD4/EFA6B* could act as a haplo-insufficient tumor-antagonist gene. Cumulative evidence currently highlights the role of heterozygosity in tumor progression^54,55^. In BC, mutations and copy number alterations of *PSD4/EFA6B* gene, located in chromosome 2q14, are very rare^7,34^. It therefore seems more likely that the loss of EFA6B expression observed in BC patients is due to events affecting its messenger and/or protein abundance^53^.

There have been no genomic studies showing *PSD4/*EFA6B to be a tumor suppressor gene and our orthotopic xenograft experiments of MCF10A EFA6B KO cells in immunosuppressed mice did not produce any tumor arguing that the loss of EFA6B alone is not a driver mutation. Yet, non-transforming secondary driver mutations provide selective advantages to transformed cells^56^. The discovery that the simple loss of EFA6B can model cancer invasion implies that such alteration can orchestrate many of the steps of the invasion cascade independently of an oncogenic mutation, congruent with the two-hit hypothesis, one that will transform the cell and another one that will help overcome the control by the microenvironment^57^. Thus, the loss of expression of EFA6B could act as a secondary driver mutation that would confer invasive properties and high metastatic susceptibility to transformed cells. Consistent with this hypothesis, we have shown here that invasive BCs had a diminished expression of *PSD4* when compared to pre-invasive *in situ* BCs and that EFA6B KO cells have a transcriptomic signature that shares several gene ontologies with a signature of progression from DCIS to IDC. Furthermore, *PSD4* expression also decreased from the primary BC to the metastatic sample in a series of 29 matched pairs, and *PSD4* down-regulation was associated with poor-prognosis features and shorter DFS in a large series of non-metastatic BC. These are key observations with respect to the treatment of patients diagnosed with DCIS and early invasive BC^58^. Further studies of the novel EFA6B-regulated pathway could help to better understand, predict and treat the progression of DCIS towards invasive BC and of invasive BC towards the lethal metastatic disease.

## Acknowledgements

This work was supported by the Centre National de la Recherche Scientifique (CNRS) and the Fondation Association pour la Recherche sur le Cancer (ARC). RF was supported by the Agence Nationale pour la Recherche (ANR) through the Investissement pour le Futur, Labex Signalife Program ANR-11Labx-0028-01. MVR was supported by a CONACYT fellowship from the Mexican government. PF, ML, DB and FB are supported by the Ligue Nationale Contre le Cancer (label) and Fondation Groupe EDF. We would like to thank Pr. Robert A. Weinberg (Whitehead Institute for Biomedical Research, MA, USA), for providing the HMLE cells and Dr. P. Chavrier (Institut Curie, Paris, France) for sharing his expertise. The results shown here are in part based upon data generated by the TCGA.

## Supplementary Figure legends

**Supplementary Figure 1**: EFA6B knock-out does not affect cell proliferation nor migration but stimulates the formation of degradative invadopodia. **a**) MCF10A WT and EFA6B KO55 cells were plated at 5×10^5^ cells per well in a 12-well dish and counted over 5 days using the LUNA™ cell counter (Logos Biosystems). The graph represents the mean of 3 independent experiments performed in triplicates ±SEM. **b**) MCF10A WT and EFA6B KO55 cells were plated at confluency in a 12-well dish (10^6^ cells per well). The next day a wound was performed using a pipet tip. After a wash with complete medium, the closure of the wound was followed overtime by phase contrast video-microscopy using a Cytation5 (Biotek). The images taken every 15min were processed and the area of the wound closure calculated using the software ImageJ. The graph represents the percentage of closure relative to the surface area of the wound at t=0 of 6 independent experiments performed in triplicates ±SEM. **c**) EFA6B KO55 cells were stained for F-actin (red) and cortactin (blue) 48h after plating on Oregon488-gelatin (green) coated coverslips. The left image is the Oregon488-gelatin staining to visualize the digested spots. The right image is the merge of all three markers to visualize the co-localization of cortactin with F-actin, appearing in purple, coincidental with the spots of digested gelatin. Scale bars 10 μm. The graph shows the fluorescent intensity of all three markers over the indicated line scan on the image. Arrowheads point to three invadopodia selected along the line scan.

**Supplementary Figure 2**: Gene set enrichment analysis (GSEA) of *PSD4* KO *versus* WT MCF10A cells, showing enrichment in the WT cells of the “Matrisome+receptors” gene set (KEGG_FA_ECM, including genes of the KEGG gene sets Focal Adhesion (‘FA’), ECM-receptor interaction (‘ECM’) and the human matrisome database) (**a**), and of the Gene Ontology “cell-cell adhesion” signature (**b**).

**Supplementary Figure 3**: EFA6B knock-out induces EMT without the acquisition of stemness characteristics. **a**) Representative images of the indicated spheroids grown 2 days in collagen I stained for E-cadherin or N-cadherin. Scale bars 20μm. **b**) Representative bright field phase contrast images of the indicated cells grown 2 days in hanging drops, n=6. **c**) The MCF10A WT (blue) and EFA6B KO55 (green) cells were labeled for the cell surface markers EpCAM and CD49f. The red dotted line circles a new de-differentiated population EpCAM^-/low^ and CD49f^low^ (11.7%) appearing in the EFA6B KO55 cell line. **d**) Expression of EMT-associated genes by qPCR analysis in HMLE Het.25 cells normalized to HMLE WT23. **e**) Expression of the indicated EMT-TF genes by qPCR analysis 2 days post-transfection in MCF10A EFA6B KO55, HMLE EFA6B KO3 and KO1 cells with the indicated corresponding specific siRNAs normalized to siCtl. n=5, average ±SEM. **f**) Percentage of ALDH1 positive cells in MCF10A WT and EFA6B KO55 cell lines analyzed by FACS. **g**) Percentage of CD44^+^/CD24^-/low^ cells in MCF10A WT and EFA6B KO55 cell lines analyzed by FACS. **h**) The cell viability of both MCF10A WT and EFA6B KO55 cells after three days of drug exposure at the indicated concentrations was determined by alamar blue-based assay (Invitrogen). Fluorescence intensity was measured at 530 nm and 590 nm wavelength for excitation and emission using a microplate reader (Infinite® M1000 Pro, TECAN). The graph curves represent the mean percentage of viability relative to the respective untreated controls of three independent experiments ±SEM.

**Supplementary Figure 4**: EFA6B KO2 cells display contractility and invasive properties similar to the KO55 cells. **a**) Expression of the indicated proteins analyzed by immunoblot in MCF10A WT, EFA6B KO55 and EFA6B KO2 cells. **b**) Representative images of the EFA6B KO 2 cells grown 2 days on collagen I-coated coverslips and stained for cortactin (green), F-actin (red) and nuclei (blue). The large inset is a 2X zoom-in image of the indicated area. Scale bar 20 μm. **c**) Quantification of the percentage of indicated cells displaying invadopodia. **d**) Quantification of the contractility of EFA6B KO2 cells transfected with the indicated siRNA evaluated by a collagen gel contraction assay. Values are mean surface area of the collagen gel ±SEM. n*=3*, Student’s *t*-test *p* values were * *p < 0.05*; *** *p < 0.001*. **e**) Quantification of the percentage of aggregates with invasive protrusions of EFA6B KO2 cells grown in collagen I for 2 days post-transfection with the indicated siRNAs. n=3, average ±SEM, Student’s *t*-test *p* values were **** *p < 0.0001*; *** *p < 0.001*. **f**) Quantification of the percentage of invadopodia in EFA6B KO2 cells grown in collagen I for 2 days post-transfection with CDC42 targeted siRNAs. n=3, average ±SEM, Student’s *t*-test *p* values were ** *p < 0.01*. **g**) Representative images of EFA6B KO 2 cells transfected with the indicated siRNAs and stained for cortactin (green), F-actin (red) and nuclei (blue). Scale bars 20μm.

## Supplementary Tables

**Supplementary Table 1:** List of 296 genes differentially expressed between the *PSD4* KO MCF10A cells (N=5, including KO55 (N=3) and KO2.9 cells (N=2)) and the *PSD4* WT MCF10A cells (N=3)

**Supplementary Table 2:** Gene ontologies of the 296 genes differentially expressed between the *PSD4* KO MCF10A cells (N=5, including KO55 (N=3) and KO2.9 cells (N=2)) and the *PSD4* WT MCF10A cells (N=3)

**Supplementary Table 3:** List of 34 BC data sets included in the study.

**Supplementary Table 4:** Correlations of *PSD4* expression with clinico-pathological features of BC.

**Supplementary Table 5:** List of primary antibodies

**Supplementary Table 6:** List and sequence of RTqPCR oligonucleotides

## Methods

### Cells and antibodies

MCF10A cells were obtained from ATCC (LGC Standards, France) and grown in DMEM/F-12 (1:1), horse serum 5%, non-essential amino acids 1%, insulin 10μg/ml, hydrocortisone 1μg/ml, EGF 10ng/ml, cholera toxin 100ng/ml and penicillin (100u/ml)-streptomycin (100μg/ml). HMLE cells were obtained from Dr. Robert A. Weinberg (Whitehead Institute for Biomedical Research, Cambridge, MA, USA)^59^ and grown in DMEM/F-12 (1:1), fetal calf serum 10%, insulin 10μg/ml, hydrocortisone 0.5μg/ml and penicillin (100u/ml)-streptomycin (100 μg/ml). All culture reagents were from Invitrogen (Fisher Scientific, France) except for the fetal calf serum (Dutscher, France). Unless otherwise indicated, all other reagents were from (Sigma-Aldrich, France). All secondary antibodies and fluorescent probes were from Molecular Probes (Invitrogen). For the list of primary antibodies refer to the Supplementary Table 5.

### 3D culture

3D culture was performed using rat tail Collagen I (Corning^®^, Fisher Scientific, France). The collagen I solution was neutralized using 1N NaOH and diluted in PBS to a final concentration of 1mg/ml for the contractility assay, and 2mg/ml for the invasion assay.

### CRISPR/Cas9 knock-out

The *PSD4* KO mutation was obtained using the CRISPR/Cas9 gene editing technology^60^. Two guided RNAs targeting the exon 1 of human *PSD4* were selected from the crispr.mit.edu/ and crispor.tefor.net/ websites. Off-target specificity was assessed by BLAST analysis of the human genome. The guided RNAs were cloned separately into the vector pSpCas9(BB)-2A-GFP (PX458) encoding for Cas9 and GFP (Addgene plasmid #48138). Sequences were as follows: PSD4-Z1T8-pX458-FOR1 forward 5’ CACCgaggatccaccggagcctttcg 3’, PSD4-Z1T8-pX458-REV1 reverse 5’ AAACcgaaaggctccggtggatcctc 3’, PSD4-Z2T36-pX458-FOR2 forward 5’ CACCgttctctgagcaaggactcgcc 3’, PSD4-Z2T36-pX458-REV2 reverse 5’ AAACggcgagtccttgctcagagaac 3’. HMLE and MCF10A cells were transfected using Lipofectamin 3000 (Invitrogen) according to manufacturer’s recommendations. 24h post-transfection, GFP positive cells were sorted and cloned in 96-well culture plates using the BD FACSAria III (BD Biosciences). The isolated clones were screened by immunoblot and the mutations validated using the Surveyor^®^ assay (Integrated DNA Technologies) followed by genomic sequencing of the targeted sequence. The three different populations of HMLE were sorted according to their expression of EpCAM and CD49f as shown in Figure 2 before next-day transfection.

### RNA isolation and RT-qPCR

Total RNA was isolated using the Tri Reagent (Sigma) and treated with Ambion™ Dnase I (Invitrogen) following the manufacturers’ instructions. RNA quality was tested using an Agilent Bioanalyzer. 2 μg of total RNA was denatured at 65°C for 10 min and incubated for 1h at 50°C in the presence of 2.5 mM dNTP, 100 U Superscript III (Invitrogen) using 0.5 μg oligo(dT)15 primer in a total volume of 20 μl, followed by an inactivation step of 15 min at 70°C. A control PCR reaction of the reverse transcription was performed with human GAPDH forward (gaacatcatccctgcatcc) and reverse (ccagtgagcttcccgttca) primers with Q5 High fidelity DNA polymerase according to the standard protocol described by the manufacturer (New England BioLabs^®^). The PCR products obtained after 35 cycles were separated through a 1% agarose gel, visualized under UV after staining with ethidium bromide. Real-time PCR was carried out with The LightCycler^®^ 480 SYBR Green I Master (Roche Life Science) in triplicates and analyzed using LightCycler^®^ 480 Software, Version 1.5 (Roche). The expression of each gene was normalized to the HPRT1 or GAPDH housekeeping gene and relative levels were calculated on the basis of the comparative cycle threshold Ct method (2-ΔΔCt) where ΔΔCt is the difference in Ct between target and reference gene. For the list and sequence of RTqPCR oligonucleotides refer to the Supplementary Table 6.

### Transcriptomic analyses

DNA-microarrays were used to define and compare the transcriptional profiles of MCF10A harboring *PSD4* homozygous (i.e. KO55, N=3) or heterozygous (i.e. Het2.9, N=2) knock-out and WT MCF10A as control (N=2). Experiments were done as recommended by the manufacturer (Affymetrix, Thermo Fisher) from 100 ng of total RNA for each sample using the Affymetrix GeneChip HuGene 2.0 ST arrays. Expression data were normalized by RMA with the non-parametric quantile algorithm in R using Bioconductor and associated packages (version 3.5.2; http://www.cran.r-project.org/). Before analysis, expression data were filtered to remove probes with low and poorly measured expression and standard deviation inferior to 0.25 log2 units across samples, resulting in 5,640 genes. We compared the expression profiles between the MCF10A harboring *PSD4* knock-out (N=5) and the MCF10A control (N=2) using a moderated t-test with empirical Bayes statistic^61^ included in the limma R packages (version 3.38.3). False discovery rate (FDR)^62^ was applied to correct the multiple-testing hypothesis. Significantly differentially expressed genes were defined by the following thresholds: p-value<0.05, q-value<0.15 and fold change FC>|2x|. Ontology analysis of the resulting gene list was based on the Gene Ontology (GO) terms using the Database for Annotation, Visualization and Integrated Discovery (DAVID; david.abcc.ncifcrf.gov/). Furthermore, to explore the impact of *PSD4* knock-out in MCF10A on the matrisome and its receptors, we applied Gene Set Enrichment Analysis (GSEA) (http://www.broadinstitute.org/gsea/) for comparing the expression profiles of *PSD4* KO *versus PSD4* WT MCF10A cells. GSEA was based on two gene sets define by the GO terms list ‘Cell-cell adhesion’ and a gathering of the KEGG gene sets Focal Adhesion (‘FA’, hsa04510), ECM-receptor interaction (‘ECM’, hsa04512) and the human matrisome database (version August 2014; Hynes Lab; Naba A, Ding H, Whittaker CA, Hynes RO. http://matrisomeproject.mit.edu) defining our “Matrisome+receptors” gene set. We used the class differential metric for ranking these filtered genes, weighted enrichment statistic for computing enrichment score (ES) of each gene set tested and 1000 permutations to evaluate significance as parameters for the GSEA.

To determine *PSD4* mRNA expression in clinical BC samples, we collected and analyzed publicly available transcriptomic data. First, we compared the expression of the four *PSD* genes in a series of 53 ductal carcinomas *in situ* (DCIS) and 51 invasive ductal carcinomas (IDC), including 11 cases for which separate samples of tumor epithelium and adjacent stroma had been generated by laser-capture micro-dissection (LCM) before profiling^31^. Expression profiling was based on DNA microarrays and comparison of expression levels between DCIS and IDC was done using the Student’s *t*-test. Second, we analyzed expression from 29 matched metastasis-primary BC pairs, collected through two data sets: TCGA^34^ for 7 pairs, and Vareslija’s study^35^ for 22 pairs, all having been profiled using RNA-seq. The comparison of *PSD4* expression levels between metastases and primary BC was done using the paired Mann-Whitney test. Finally, in order to search for correlations with clinico-pathological parameters in invasive BC, we analyzed DNA microarray- and RNA-seq-based data generated across 34 retrospective data sets including ours (**Supplementary Table 3**). This resulted in a total of 8,464 non-redundant primary invasive BC and four normal breast (NB) samples. Analysis was done as previously reported^63^. PSD4 down-regulation was defined by a ratio tumor/NB ≤0.5, and no down-deregulation by a ratio >0.5. To avoid biases related to immuno-histochemical and pathological analyses across different sets and to increase the amount of available data, ER and ERBB2 statutes were defined as positive or negative using respective mRNA expression data of *ESR1* and *ERBB2* as described previously^7^. To compare the distribution according to categorical variables, we used the correlation of *PSD4* expression (down *versus* no down) with the clinico-pathological features using the Student’s t-test for continuous features and Fisher’s exact test for discrete features. Disease-free survival (DFS) was calculated from the date of diagnosis until date of relapse or death. Follow-up was measured from the date of diagnosis to the date of last news for patients without event. Survivals were calculated using the Kaplan-Meier method and curves were compared with the log-rank test.

### Transient transfection

Transient transfection of siRNAs was performed using RNAiMAX (Invitrogen) according to the manufacturer instructions. All siRNAs were from Dharmacon ON-TARGETplus collection (Horizon Discovery).

### Stable transfection

HEK-293T cells were transfected with the human *PSD4* containing pLV lentivirus (VB160428-1095xdp – Vector Builder) together with the 3^rd^ generation lentiviral packaging plasmid (Addgene): pMDLG/pRRE (Gag and Pol), pRSV-Rev (Rev), and pMD2.G (VSV-G envelope) using JETPEI (Polypus Transfection). Supernatants containing the lentiviruses were collected at 48h and 72h post-transfection. MCF10A cells were transduced with the filtered supernatants in the presence of 10 μg/ml of Polybrene (Sigma-Aldrich). Transduced cells were selected with 250 μg/ml of G418 (Sigma).

### Immunoblot

Cells grown on plastic dishes were washed 3 times with phosphate buffered saline (PBS) and lysed with an SDS lysis buffer containing 0.5% SDS, 150 mM NaCl, 5 mM EDTA, 20 mM Triethanolamine–HCl pH 8.1, 1 mM PMSF. The lysate was heated at 95°C for 10 min and then thoroughly vortexed for 15 min. After centrifugation at 16,000 *g* for 20 min at room temperature, the supernatant was transferred into a new tube containing 5 × Laemmli buffer and further boiled 5min. For protein analysis, cell lysates were loaded into SDS-PAGE and transferred onto a nitrocellulose membrane. Membrane blocking and secondary antibodies dilutions were done in PBS 5% non-fat dry milk, primary antibodies were diluted in PBS containing 3% BSA. The proteins were revealed by chemiluminescence (ECL™, Amersham France) using secondary antibodies directly coupled to HRP. The membranes were analyzed with the luminescent image analyzer Fusion (VILBER, France).

### Flow cytometry

Cells were detached using Accutase (Stemcell technologies, NC, USA) and washed 3 times in PFE (PBS, 2mM EDTA, 2% Foetal Bovine Serum). The cells were incubated at 4°C for 30 min in the presence of the indicated antibodies diluted in PFE. After washes, cells resuspended in cold PFE were examined by BD LSRII Fortessa™ cell analyser. The ALDEFLUOR kit (Stemcell Technologies) was used to quantify the ALDH enzymatic activity. Results were processed using Kaluza or FlowJo softwares.

### Invasion assay

Cells transfected with siRNA were plated in collagen 24h after transfection. Human mammary individual cells (100 cells/μl) resuspended in 200 μl of collagen (2mg/ml) were plated in chambered cover glass (Lab-Tek, Fisher Scientific), and incubated for 30 min at 37°C in 5% CO2 before addition of complete medium. After 48h, protrusions were quantified using a phase contrast microscope (Leica) by counting 100 cellular aggregates per well. Cells with at least one membrane extension of at least 2 microns’ length were considered invasive.

### Fluorescent gelatin degradation assay

Oregon Green 488-labeled gelatin was from Molecular Probes (Invitrogen). Sterilized coverslips (18-mm diameter) were coated with 50 μg/ml poly-L-lysine for 20 min at RT, washed with PBS, and fixed with 0.5% glutaraldehyde for 10 minutes. After 3 washes with PBS coverslips were inverted on a 40-μl drop of 0.2% fluorescently labeled gelatin in 2% sucrose in PBS and incubated for 10 min at RT. After washing with PBS, coverslips were incubated in a 5 mg/ml solution of borohydride for 3 min, washed three times in PBS, and incubated with 1 ml of complete medium for 30 minutes. 5 x 10^4^ cells per 12-well were plated on the fluorescent gelatin-coated coverslips and incubated at 37°C for 48 h. Cells were then washed three times with PBS and fixed with 4% PFA for 20 min and processed for labeling with Texas Red-Phalloidin and DAPI. The coverslips were mounted with Mowiol mounting medium. Cells were imaged on a confocal microscope (Leica TCS-SP5). For the quantification of the gelatin degradation, the total area of degraded matrix measured with ImageJ software was divided by the total area of each phalloidin-labeled cell as described elsewhere^64^. A total of 90 cells were analyzed in three independent experiments.

### Quantification of cellular collagenolysis

2 × 10^4^ MCF10A cells were re-suspended in 0.2 ml of 2 mg/ml collagen I solution loaded on a Lab-Tek 8-well chambered cover-glass. After gelling for 1h at 37°C, a complete DMEM-F12 medium was added, and collagen-embedded cells were incubated 48h at 37°C in 1% CO_2_. After fixation in 4% PFA for 20 min, samples were incubated with collagen type I cleavage site antibody (Col1-2/3 C) diluted 1:200 in PBS (5 μg/ml, 24h at 4°C), washed extensively with PBS, and counterstained with Cy5-conjugated anti-rabbit IgG antibodies, DAPI and Texas Red-phalloidin. Images acquisition was performed with a confocal microscope (Leica TCS-SP5) with a 60x oil objective. Quantification of degradation spots was performed with a homemade ImageJ program as described elsewhere^64^. The degradation index is the number of degradation spots divided by the number of cells in each cluster present in the field. A total of 90 cells were analyzed in 3 independent experiments.

### Immunofluorescence

Cells were fixed in 4% paraformaldehyde for 20 min, washed three times in PBS and permeabilized in PBS containing 0.05% saponin, 0.5% horse serum for 30 min at 37°C. Then the cells were incubated with primary antibodies over-night and counterstained with appropriate fluorescent secondary antibodies, DAPI or phalloidin. Images acquisition was performed with a confocal microscope (Leica TCS-SP5) with a 60x oil objective.

### Confocal reflectance microscopy

MCF10A spheroids were embedded in collagen I for 48h at 37°C, then spheroids were fixed, and immunofluorescence was performed as indicated above. The imaging of the orientation of the collagen matrix fibers was performed by confocal reflectance microscopy using a scanning confocal microscope (Leica TCS-SP5). The collagen gels were excited with a 488 nm laser, and signal between 485 nm and 495 nm was collected.

### Contractility assay

70 μl of collagen mixed with human mammary cells (700 cells/μl) were plated in 96-well plates, and incubated for 30 min at 37°C in 5% CO2 before addition of complete medium. The collagen gels were detached from the plastic using a pipet cone. Collagen area was quantified using Image J software.

### Pull down assay

MCF10A cells were lysed at 4°C in 0.5% Nonidet P-40, 20 mM Hepes pH 7.4, 125 mM NaCl, 1 mM phenylmethylsulphonyl fluoride (PMSF) and a cocktail of protease inhibitors (Complete, Roche Diagnostics). The cleared lysates containing the protein of interest were incubated 4h at 4°C with 1.5 μM of the indicated GST-fused proteins and 30 μl of glutathione-sepharose CL-4B beads (GE Healthcare). After three washes in lysis buffer, the beads were boiled in Laemmli buffer, submitted to SDS–PAGE and the indicated proteins were revealed by immunoblot. GST fused to the ARF6GTP binding domain of ARHGAP10, the CDC42/RAC binding domain of PAK, or the RHO binding domain of Rhotekin were used to pull-down ARF6GTP, CDC42GTP/RAC1GTP or RHOAGTP, respectively.

### Integrin antibody blocking assay

MCF10A cells were trypsinized, washed with PBS and re-suspended in complete DMEM-F12 medium. The cells were then incubated for 30 min at 37°C to allow for recovery of cell surface receptors. 5 x 10^4^ cells were re-suspended in 0.2 μl of a 2 mg/ml collagen I solution and control or blocking antibodies were added. The cell suspension was loaded on an 8-well Lab-tek chambered cover-glass and left to gelify for 1h at 37°C. Complete DMEM-F12 medium containing the control or blocking antibodies was added on top of collagen-embedded cells. The samples were incubated 48 h at 37°C in 1% CO2. The development of protrusions was quantified as described above by counting 100 cell clusters, in 3 independent experiments.

### Statistics

Experiments were performed at least three times independently. The number of repeats is indicated in the legend as (n). Error bars represent ± standard error of the mean (SEM). Statistical significance was determined using a Student *t*-test and one-way ANOVA, in which *p* values < 0.05 were considered statistically significant. All statistical analyses have been performed with GraphPad Prism software.

