## Supplementary figures and images for "EFA6B regulates a stop signal for collective invasion in breast cancer"

### Supp Figures

#### Supplementary Figure 1

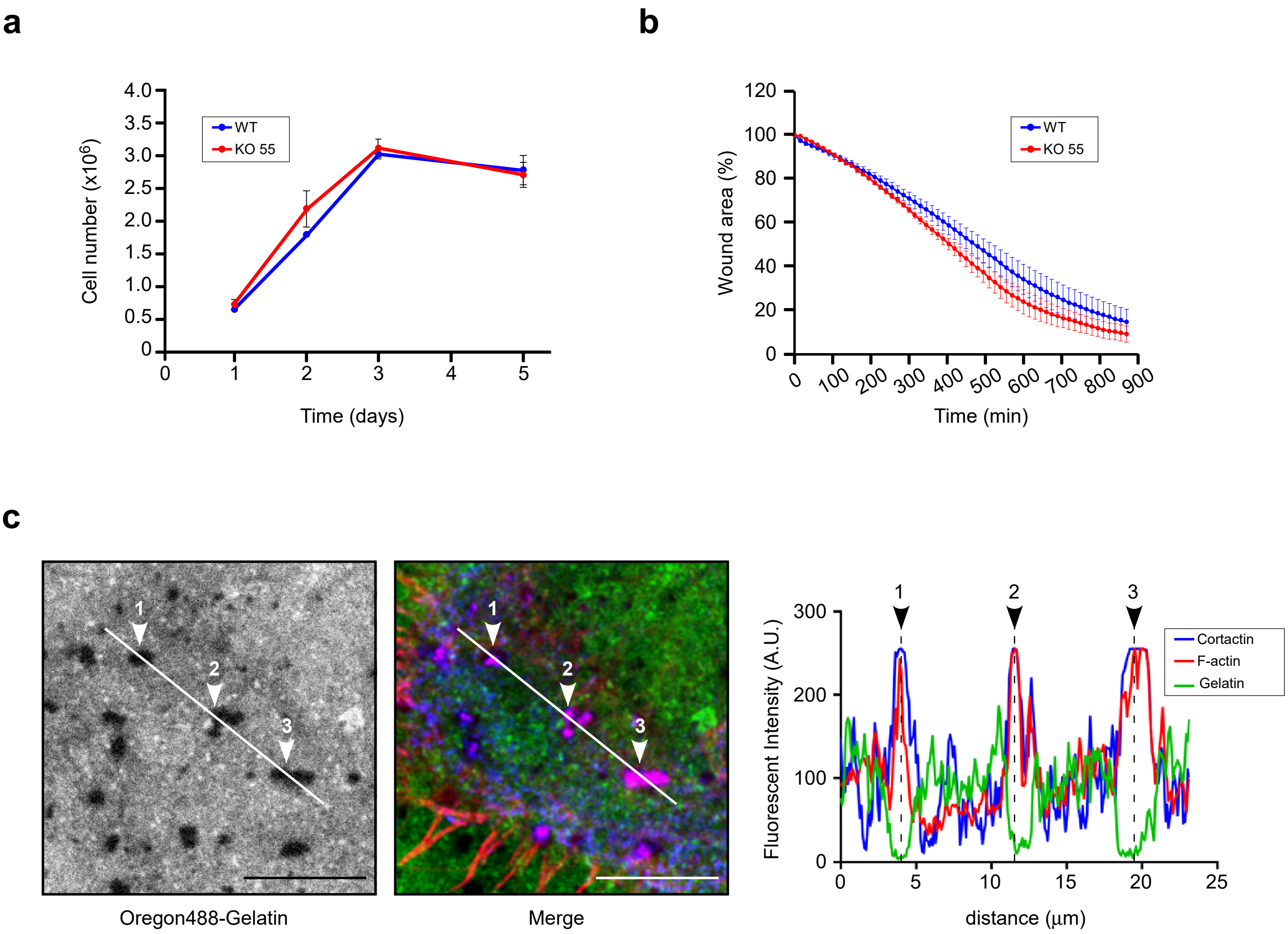

#### Supplementary Figure 2

a

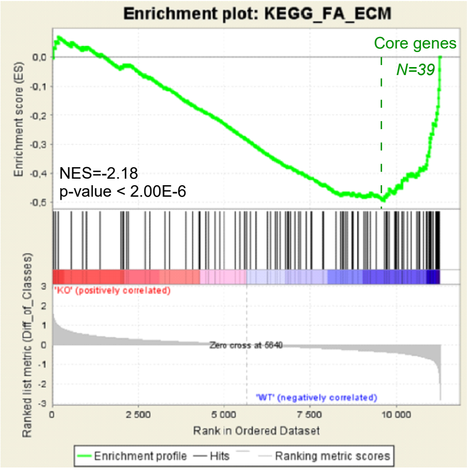


b

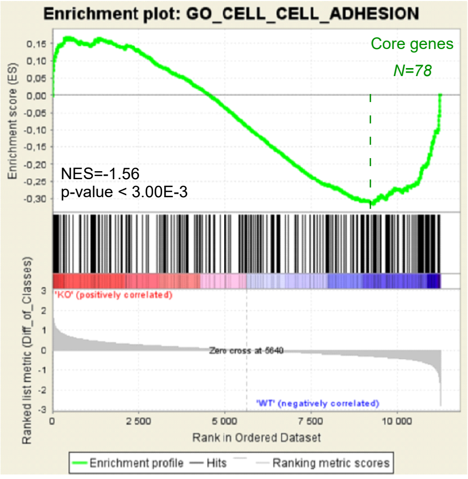

#### Supplementary Figure 3

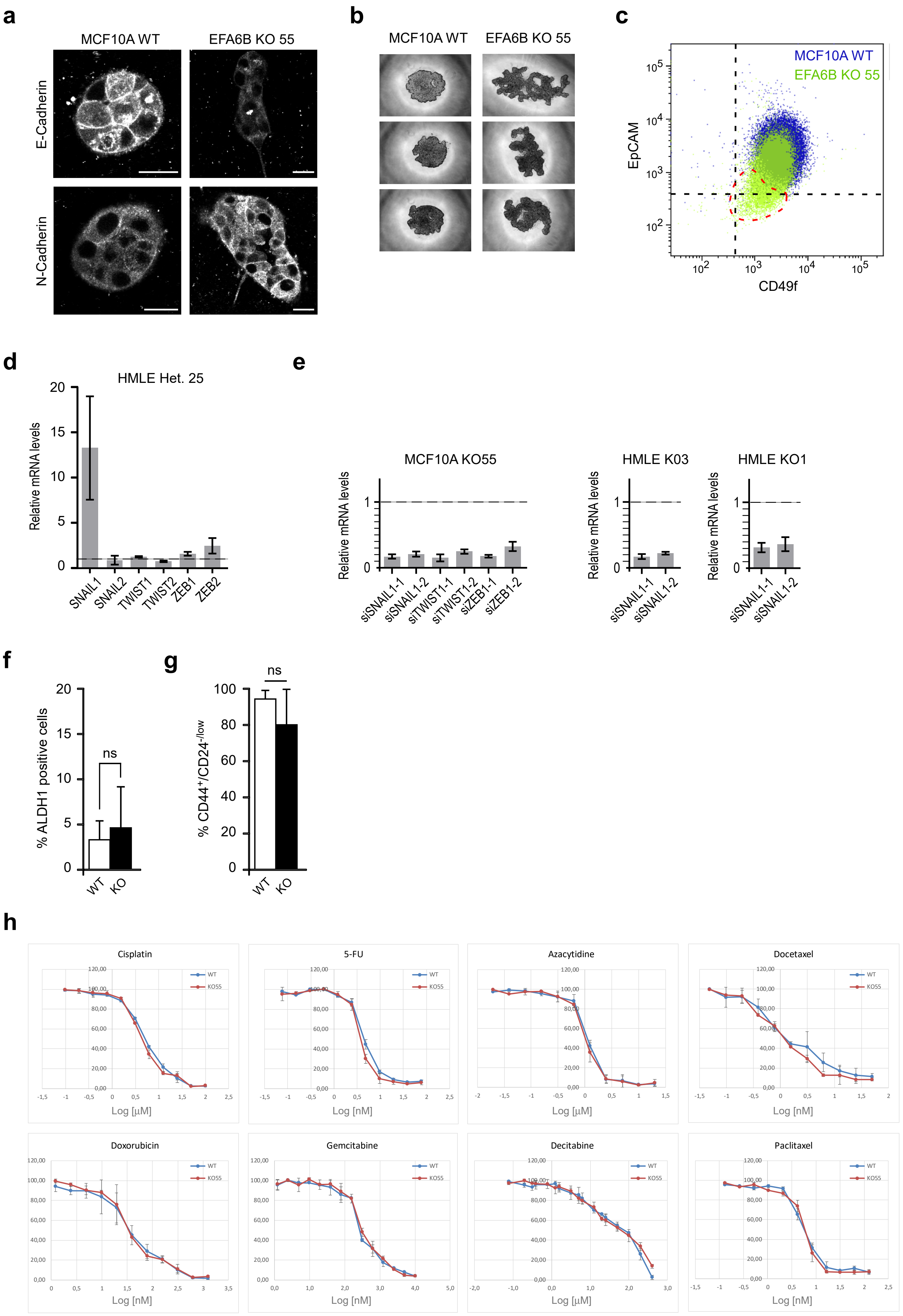

#### Supplementary Figure 4

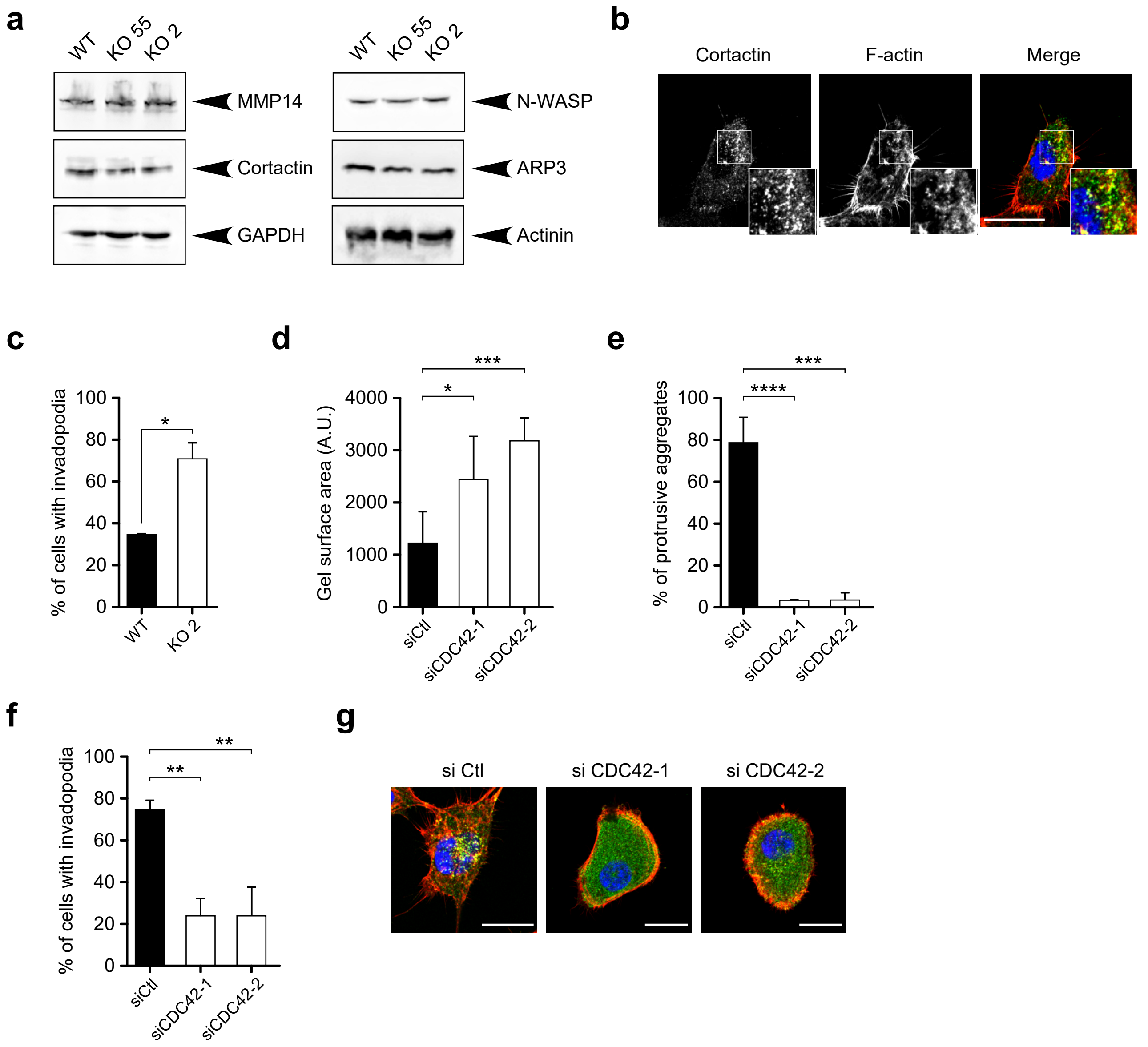
