## Supplementary Table S5 for "EFA6B regulates a stop signal for collective invasion in breast cancer"

| **Antigen** | **Antibody source and reference** | **Application** |
| --- | --- | --- |
| Integrin-β1 | Santa Cruz, sc-53711 | Immunoblot |
| Integrin-β1 | Santa Cruz, sc-13590 | Flow cytometry Immunofluorescence  Blocking antibody |
| Integrin-β4 | BD Biosciences, 611232 | Immunoblot Immunofluorescence |
| Integrin-β4 | BD Biosciences, 555719 | Flow cytometry |
| Integrin-β4 | Millipore, MAB2059 | Blocking antibody |
| Integrin-α2 | Santa Cruz, sc-53502 | Flow cytometry Immunofluorescence |
| Integrin-α2 | Millipore, MAB1950 | Blocking antibody |
| Integrin-α3 | Millipore, MAB2057 | Flow cytometry Immunofluorescence Blocking antibody |
| Integrin-α6 | BD Biosciences, 555734 | Flow cytometry Immunofluorescence Blocking antibody |
| EpCAM | BD Biosciences, 563181 | Flow cytometry |
| CD49f | BD Biosciences, 562582 | Flow cytometry |
| CD24 | BD Biosciences, 561644 | Flow cytometry |
| CD44 | BD Biosciences, 560568 | Flow cytometry |
| N-cadherin | BD Biosciences, 610921 | Immunofluorescence Immunoblot |
| E-cadherin | Thermo Scientific, 33-4000 | Immunofluorescence |
| E-cadherin | BD Biosciences, 610182 | Immunoblot |
| Vimentin | Sigma, V6389 | Immunoblot |
| Claudin 1 | ZYMED Laboratories, 51-9000 | Immunoblot |
| Claudin 3 | Millipore, 2819163 | Immunoblot |
| Occludin | Thermo Scientific, 40-4700 | Immunoblot |
| ARF1 | Novus Biologicals, NB100-55421 | Immunoblot |
| ARF5 | Abnova, H00000381-M01 | Immunoblot |
| ARF6 | Gift from Dr. Bourgoin | Immunoblot |
| EFA6A | Gift from Dr. Sakagami | Immunoblot |
| EFA6B | Sigma, HPA034722 | Immunoblot |
| EFA6D | Gift from Dr. Sakagami | Immunoblot |
| MMP-14 | Millipore, MAB3328 | Immunoblot |
| Cortactin | Millipore, 05-180 | Immunofluorescence  Immunoblot |
| N-WASP | Cell Signaling, 4848 | Immunoblot |
| ARP3 | BD Biosciences, 612134 | Immunoblot |
| pMLC | Cell Signaling, 3671 & 3675 | Immunoblot |
| MLC | Sigma, M4401 | Immunoblot |
| CDC42 | BD Biosciences, 610928 | Immunoblot |
| RAC1 | BD Biosciences, 610650 | Immunoblot |
| RHOA | Santa Cruz, sc-418 | Immunoblot |
| ROCK 1 | Santa Cruz, sc-17794 | Immunoblot |
| ROCK 2 | Santa Cruz, sc-398519 | Immunoblot |
| Actin | Sigma, A4700 | Immunoblot |
| Actinin | Sigma, A5044 | Immunoblot |
| GST | GE Healthcare, 27-4577-01 | Immunoblot |
| Hsp60 | Sigma, SAB4501464 | Immunoblot |
| p85 | Millipore, ABS1856 | Immunoblot |
| GAPDH | Sigma, G9545 | Immunoblot |
