## Supplementary Table S6 for "EFA6B regulates a stop signal for collective invasion in breast cancer"

| ***Oligo name*** | ***Sequence forward (5’-3’)*** | ***Sequence reverse (5’-3’)*** |
| --- | --- | --- |
| **E-Cadherin** | TGCCCAGAAAATGAAAAAGG | GTGTATGTGGCAATGCGTTC |
| **N-Cadherin** | ACAGTGGCCACCTACAAAGG | CCGAGATGGGGTTGATAATG |
| **Vimentin** | GAGAACTTTGCCGTTGAAGC | GCTTCCTGTAGGTGGCAATC |
| **CK14** | GGAAGCCGACATCAATGG | GCCTCTCAGGGCATTCATC |
| **ITGA6** | GATGCCACATATCACAAGGC | TGCATCAGAAGTAAGCCTCTCT |
| **Occludin** | TCAGGGAATATCCACCTATCACTTCAG | CATCAGCAGCAGCCATGTACTCTTCAC |
| **Claudin 7** | AGAGCACGGGGATGATGAG | CACCCATGGCTATACGGGC |
| **Snail 1** | GCTGCAGGACTCTAATCCAGA | ATCTCCGGAGGTGGGATG |
| **Snail 2** | GGGGAGAAGCCTTTTTCTTG | TCCTCATGTTTGTGCAGGAG |
| **Twist 1** | GGCTCAGCTACGCCTTCTC | CCTTCTCTGGAAACAATGACATCT |
| **Twist 2** | CCTCTGACAAGCTGAGCAAG | GCAGGACCTGGTAGAGGAAG |
| **Zeb 1** | AACTGCTGGGAGGATGACA | TCCTGCTTCATCTGCCTGA |
| **Zeb 2** | CGATCCAGACCGCAATTAAC | TGCTGACTGCATGACCATC |
| **MRCKα** | TGAATACGCCTACCGATGCT | GTGAGTCTTGCGCTTAGGTG |
| **MRCKβ** | CCGGAAGATATGGCGAGGTTC | CCATTCACGTCCAAAAGGACAT |
| **GAPDH** | TGCCTCCTGCACCACCAACT | CCCGTTCAGCTCAGGGATGA |
| **HPRT1** | TGACCTTGATTTATTTTGCATACC | CGAGCAAGACGTTCAGTCCT |
| **U1snRNA** | GGGAGATACCATGATCACGAAGGT | ATGCAGTCGAGTTTCCCACA |
